# CROssBAR: Comprehensive Resource of Biomedical Relations with Deep Learning Applications and Knowledge Graph Representations

**DOI:** 10.1101/2020.09.14.296889

**Authors:** Tunca Doğan, Heval Atas, Vishal Joshi, Ahmet Atakan, Ahmet Sureyya Rifaioglu, Esra Nalbat, Andrew Nightingale, Rabie Saidi, Vladimir Volynkin, Hermann Zellner, Rengul Cetin-Atalay, Maria Martin, Volkan Atalay

## Abstract

Systemic analysis of available large-scale biological and biomedical data is critical for developing novel and effective treatment approaches against both complex and infectious diseases. Owing to the fact that different sections of the biomedical data is produced by different organizations/institutions using various types of technologies, the data are scattered across individual computational resources, without any explicit relations/connections to each other, which greatly hinders the comprehensive multi-omics-based analysis of data. We aimed to address this issue by constructing a new biological and biomedical data resource, CROssBAR, a comprehensive system that integrates large-scale biomedical data from various resources and store them in a new NoSQL database, enrich these data with deep-learning-based prediction of relations between numerous biomedical entities, rigorously analyse the enriched data to obtain biologically meaningful modules and display them to users via easy-to-interpret, interactive and heterogenous knowledge graph (KG) representations within an open access, user-friendly and online web-service at https://crossbar.kansil.org. As a use-case study, we constructed CROssBAR COVID-19 KGs (available at: https://crossbar.kansil.org/covid_main.php) that incorporate relevant virus and host genes/proteins, interactions, pathways, phenotypes and other diseases, as well as known and completely new predicted drugs/compounds. Our COVID-19 graphs can be utilized for a systems-level evaluation of relevant virus-host protein interactions, mechanisms, phenotypic implications and potential interventions.

## 1. Main

Systemic analysis of available large-scale biological and biomedical data is critical for developing novel and effective treatment approaches. Parts of this big-data are continuously being updated and maintained by different organizations and institutions, thus, data is scattered across individual computational resources. Although these entities are biologically related and complementary to each other, the connections between data-points at different resources are not well-established and explicit at all. In addition to the connectivity problem, another issue related to biomedical data is the incompleteness in knowledge space (e.g., possible ligands of a target biomolecule, or the disease related implications of a critical mutation). There is a clear requirement for innovative computational approaches to integrate available biomedical big-data and to complete missing information with *in silico* predictions, to serve the ultimate aim of proposing novel treatment options (especially) for complex diseases.

There are numerous studies in the literature that aimed to integrate the available biomedical data^1-10^. These studies provided useful tools and methods to the life-sciences research community; however, many of them miss important functionalities that prevent them from becoming widely adopted tools/services (Supplementary Information section 1).

In this project, we aimed to address the current shortcomings by developing a comprehensive open access biomedical system entitled CROssBAR via integrating various biological databases to each other, inferring the missing relations between existing data points, and constructing informative knowledge graphs based on specific biomedical components/terms such as a disease/phenotype, biological process, gene/protein and drug/compound, or specific combinations of them. To construct the CROssBAR system, we accomplished multiple sub-projects: *(i)* the construction of the CROssBAR database to house, and its API service to serve the integrated biomedical data, *(ii)* development of deep-learning based drug/compound-target protein interaction (DTI) prediction models, *(iii) in vitro* wet-lab experiments at different levels of the project in order both to validate/assess the relevance of *in silico* generated knowledge and to contribute to the development of new treatment options for liver diseases, *(iv)* network based organization and analysis of large-scale biomedical data using knowledge graph representations, and *(v)* the establishment of an open access web-service, where users’ biomedical term queries are processed via on-the-fly generation of knowledge graphs with both tabular and network-based visualization and download options. The CROssBAR project is schematically summarized in Fig. 1a.

**Figure 1.**
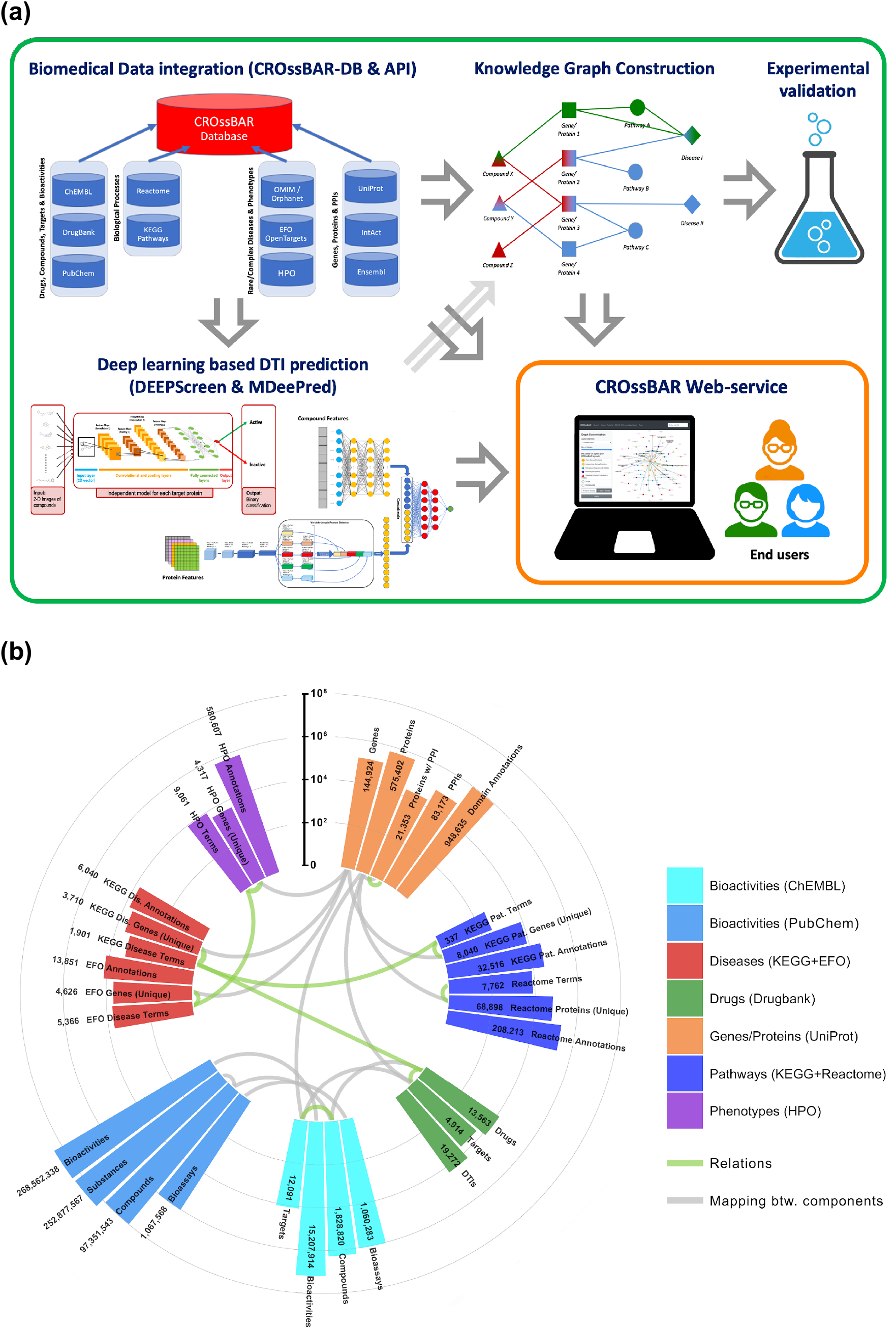
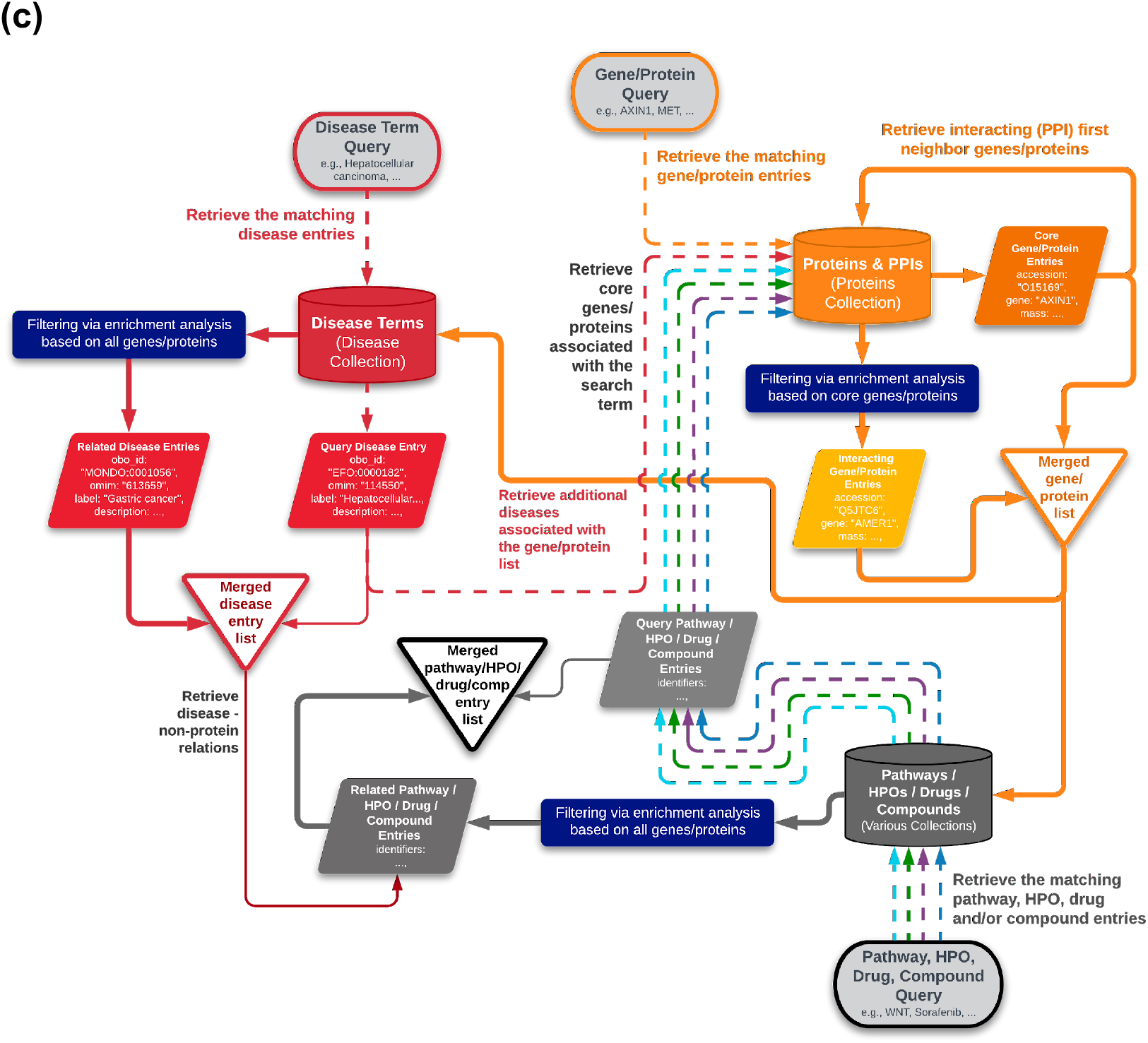
**(a)** Overall schematic representation of the CROssBAR project within five main pillars; **(b)** CROssBAR database statistics displayed in a circular bar-graph layout, bar lengths are shown in logarithmic scale, each high-level biomedical data component group is displayed with a different color, gray curves show the matching components in different database collections (i.e., bars connected with gray curves signify the same types of biomedical data -keys-, these mappings are utilized for relating independent database collections to each other), green curves signify the biomedical relations (e.g., drug-target protein interactions) between different CROssBAR components, statistics are also provided in Table S.1 (abbreviations; Dis.: disease, Pat.: pathway, DTIs: drug-target interactions); **(c)** simplified work-flow of the KG construction procedure, shown in detail for an example disease term query. With the initiation of KG construction by a disease query, the system *(1)* finds the matching disease entry from the relevant collection, *(2)* gathers genes/proteins that are associated with the query disease (i.e., core genes/proteins), *(3)* collects additional genes/proteins (i.e., first-neighbours) using PPIs of core genes/proteins, *(4)* identifies biological processes (pathways), of which these genes/proteins (core+neighbouring) are members, *(5)* gathers phenotypic terms (HPO) associated with the whole gene/protein set, *(6)* obtains known drugs and drug candidate compounds targeting these genes/proteins, together with our deep-learning based interaction predictions, and *(7)* revisits the disease collection to make another query with all collected genes/proteins, to obtain the disease entries that have similar implications as the query disease.

CROssBAR database (CROssBAR-DB) comprises carefully selected features from various data sources namely UniProt, IntAct, InterPro, Reactome, Ensembl, DrugBank, ChEMBL, PubChem, KEGG, OMIM, Orphanet, Gene Ontology, Experimental Factor Ontology (EFO) and Human Phenotype Ontology (HPO). Extract-Transform-Load (ETL) pipelines were developed for heavy lifting of data from these resources by persisting specific data attributes with the implementation of logic rules. These pipelines fetch, cleanse, validate and consolidate the data, and thus, implement a multi-omics data integration approach to release a single resource based on MongoDB collections. CROssBAR-DB, which provides a broad spectrum of information such as biomolecular functions, domains, interactions, pathways, diseases, phenotypes, drugs, compounds, etc., is hosted and maintained by the EMBL-EBI. Current data statistics of the CROssBAR-DB and the database schema are shown in Fig. 1b and in Fig. S1, respectively. CROssBAR-DB is periodically updated on demand/request basis via an automated procedure, which makes the underlying data up to date most of the time. CROssBAR-DB can be queried via a public RESTful API at: www.ebi.ac.uk/Tools/crossbar/swagger-ui.html, which provides a multi-faceted view of the stored data through 12 endpoints (Fig. S3).

As a part of the CROssBAR project, we developed two novel deep-learning-based predictive systems: DEEPScreen and MDeePred, with the aim of enriching bioactivity data by identifying unknown interactions between drugs/drug-candidate compounds and target proteins. DEEPScreen employs convolutional neural networks to process 2D structural images of drugs/compounds in 704 individually optimized high performance target-based prediction models, suited for well-studied targets^11^. MDeePred utilizes both compound and target protein features within a pairwise input hybrid deep neural network architecture to produce real valued bioactivity predictions, especially for targets with a few or no training instances^12^. We trained both systems using carefully filtered and integrated data in CROssBAR-DB, and ran our trained-models on large compound and human protein spaces to obtain comprehensive bio-interaction predictions, which are included in our knowledge graphs. We also developed an accompanying computational tool, iBioProVis, which is an unsupervised-learning-based visualization system for exploring large drug/compound-target interaction datasets in reduced dimensions^13^.

The term knowledge graph (KG) defines a specialized data representation structure, in which a collection of entities (nodes) are linked to each other (edges) in a semantic context^14^. In this study, we chose to represent heterogeneous biomedical data in KG structures. In CROssBAR knowledge graphs (CROssBAR-KG), biological components/terms (i.e., drugs/compounds, genes/proteins, bio-processes/pathways, phenotypes/diseases) are represented as nodes, and their known or predicted pairwise relationships are annotated as edges (a protein and its coding gene is treated as one merged term/entry/node). The logic behind the construction of a knowledge graph is centered around queried biological components/terms, as shown in Fig. 1c with a work-flow diagram and with an example disease term query. At each step of the process, an overrepresentation-based enrichment analysis has been performed to select the terms that are significantly associated with the growing graph, and to discard the rest. This analysis comprises a series of hypergeometric tests, based on the recorded relations in the CROssBAR database. Here, we applied a layered construction approach, always taking the genes/proteins at the centre of the enrichment analysis. Finally, additional relation types are incorporated to the graph as edges between the existing nodes (e.g., drug-disease, disease-pathway and disease-HPO), to further enrich the provided information. During the construction of graphs, terms (nodes) and their pairwise relations (edges) are directly obtained from the CROssBAR-DB. CROssBAR-KGs clearly display the direct and indirect relations between all of the terms in the graph. These intensely-processed heterogeneous biological networks are expected to aid biomedical research, especially to infer mechanisms of diseases in relation to biomolecules, systems and candidate drugs.

We developed the CROssBAR web-service (CROssBAR-WS) to make the CROssBAR-KGs available to the public in an easily interpretable and interactive way (https://crossbar.kansil.org). KGs are presented visually on web-browsers as flexible Cytoscape^15^ networks. Users can create queries with biomedical terms, individually or in combination, to obtain the relevant graph. Combinatory term query is especially critical as it provides the ability to investigate the indirect biological relationships between the terms from both the same and different biomedical components. Since there are billions of different ways to query CROssBAR, it was not feasible to pre-calculate the resulting graphs; therefore, they are set to be constructed on-the-fly, in real-time. Several options are provided to users to customize the procedure both before the search, such as the UniProt databases to be used (UniProtKB/Swiss-Prot or UniProtKB/Swiss-Prot+UniProtKB/TrEMBL), taxons to be included, and the number of terms/nodes to include from each entity type (selected from enrichment score-based ranked lists). It is also possible to display the graph using a variety of layout options, including our in-house CROssBAR-layout (Fig. 2a). Saving options let users to store the graph in different formats, including json, figure-ready snapshots and protein-centric delimited data-tables. The interactive visualization also lets users prepare a custom display by relocating the nodes/edges as desired.

**Figure 2.**
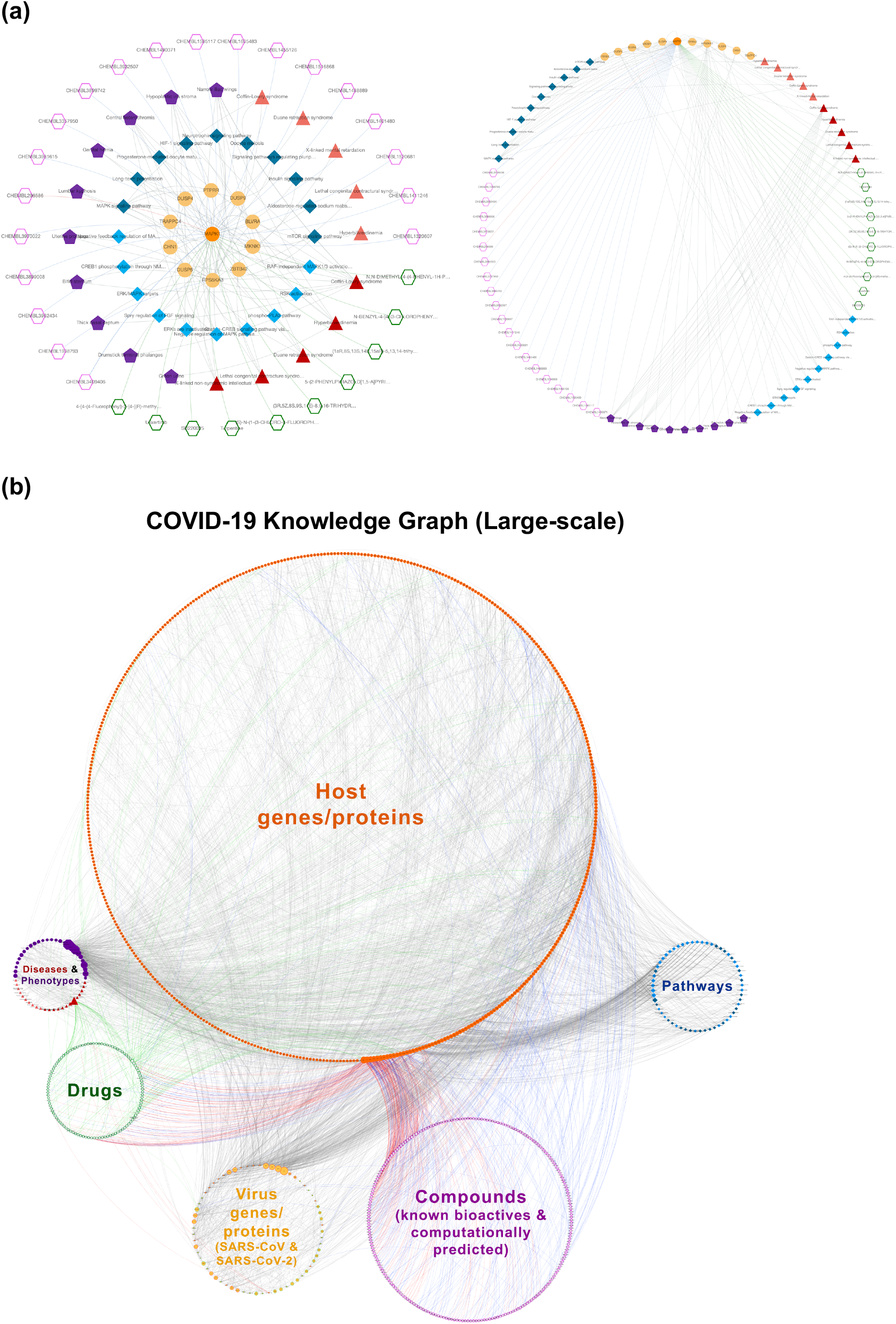
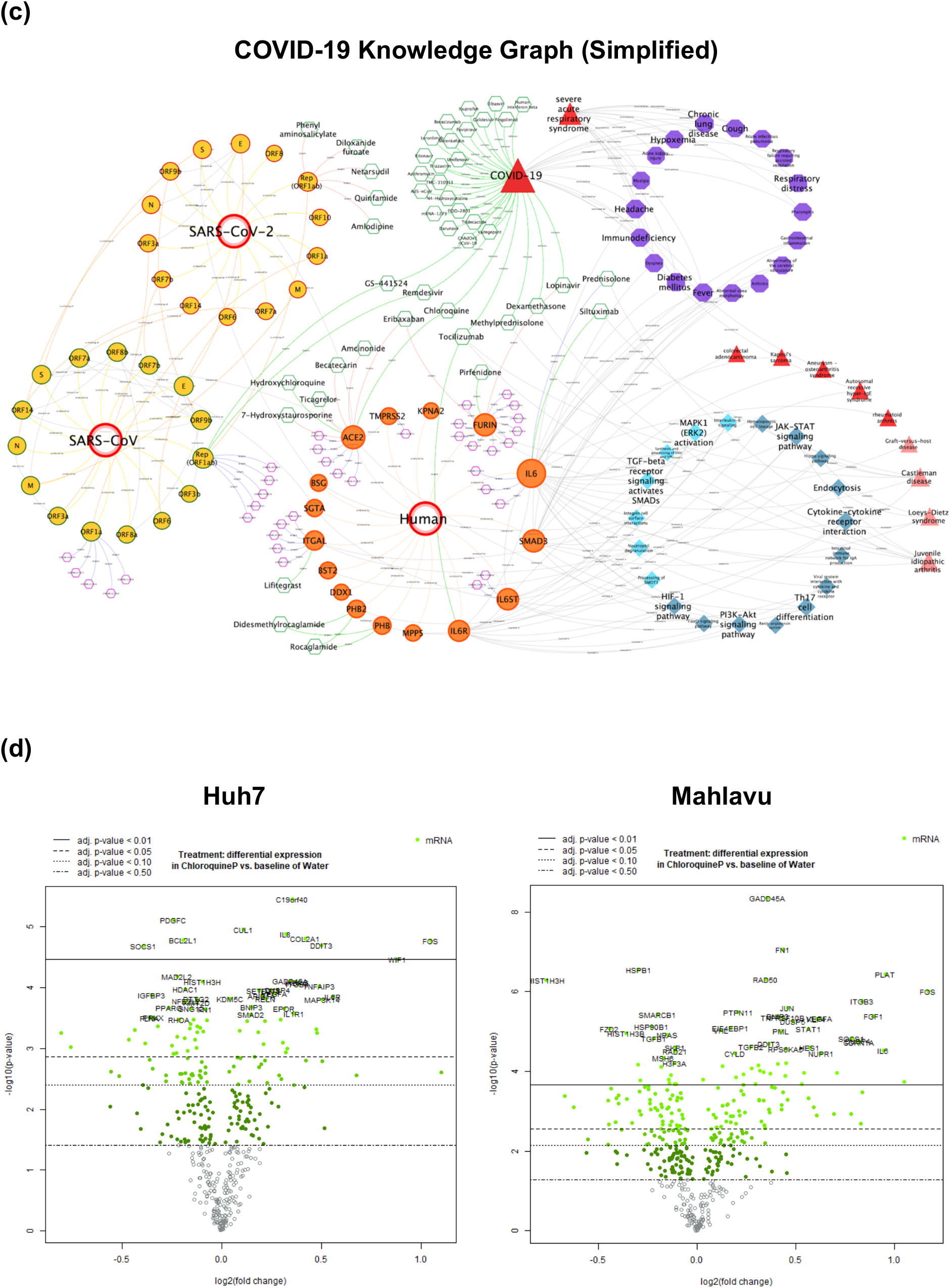
**(a)** An example KG obtained from CROssBAR-WS, generated on-the-fly with the user’s query of “MAPK1” gene (with the node limit of 10 and other default parameters), displayed under the layout selections of multi-layered CROssBAR and circular. The use case of CROssBAR COVID-19 knowledge graphs (https://crossbar.kansil.org/covid_main.php): **(b)** the large-scale KG (987 nodes and 3639 edges) and **(c)** the simplified KG (178 nodes and 298 edges). Both of these graphs reveal the most overrepresented biological processes during a SARS-CoV-2 infection (i.e., cell cycle, viral mRNA translation, endocytosis, interleukin signaling, etc.), as well as, the potential treatment options with COVID-19 related pre-clinical/clinical results (e.g., Chloroquine, Remdesivir, Favipiravir, Dexamethasone, etc.) and our novel *in silico* predictions (for both virus and host proteins) considering long-term drug discovery or short-term drug repositioning applications (e.g., Prednisolone, Tocilizumab, Isosorbide, Quercetin, Eribaxaban, Becatecarin, Quinfamide, Amlodipine, etc.). It also displays rare and complex diseases and phenotypic implications with similar host protein associations (e.g., arthritis, diabetes, respiratory distress, fever, etc.); **(d)** *in vitro* experimental results: volcano plots displaying differentially expressed genes in Chloroquine treated liver cells (Huh7 and Mahlavu). We checked the interaction between the significant DEGs (Table S.2) and genes in the large-scale COVID-19 KG, and applied Fisher’s exact test to analyse the significance of the presence of DEGs on the KG (Table S.4) as opposed to the non-DEGs in the multiplex panel of the gene expression analysis platform (NanoString). The results indicated that DEGs were significantly overrepresented (p-value < 0.01).

As a use-case, we present Coronavirus disease 2019 (COVID-19) CROssBAR-KGs (https://crossbar.kansil.org/covid_main.php). Starting from the end of 2019, the new coronavirus (SARS-CoV-2) pandemic has ravaged the entire globe and caused immeasurable damage^16^. As of July 2020, the scientific endeavour to develop effective drugs and vaccines is at peak, and systemic evaluation of the current knowledge about SARS-CoV-2 infection is expected aid researchers in this struggle. To demonstrate the capabilities of CROssBAR, we have constructed two different versions of the COVID-19 knowledge graph, (i) a large-scale version including nearly the entirety of the related information on different CROssBAR-integrated data sources, which is ideal for further network and machine learning based analysis or a detailed inspection (Fig. 2b), and (ii) a simplified version distilled to include only the most relevant genes/proteins as provided in UniProt-COVID-19 portal (https://covid-19.uniprot.org), which is ideal for fast interpretation (Fig. 2c). It is interesting to observe the indirect relations between the diseases/phenotypes in the KGs and COVID-19 over the incorporated host proteins and enriched pathways, and between COVID-19 and our in silico predicted drugs, as they may reveal further evidence to be utilized against COVID-19 (Fig. 2b,c). For this, we conducted a short literature-based validation study and found that many of these drugs have already been experimented at both preclinical and clinical stages for new COVID-19 treatments (Supplementary Information section 2).

Although COVID-19 is a respiratory disease and lung lesions have been considered the major damage caused by SARS-CoV-2, liver injury has also been reported in about one-third of hospitalized patients infected with the virus and the majority of COVID-19 patient deaths are associated with cytokine storm/release syndrome resulting in multi organ damage^17^. Hence, with the aim of indicating the biological relevance of the information in CROssBAR-KGs, we conducted *in vitro* experimentation on drug treated liver cancer cell-lines and comparatively analysed the results on both COVID-19 KGs. Chloroquine (CQ) phosphate was reported to be used in treatment of COVID-19 with controversies^18^. CQ is an anti-inflammatory drug that has been used in autoimmune diseases and can significantly alter the production of pro-inflammatory and anti-inflammatory cytokines. We investigated the effect CQ on normal hepatocytes like Huh7 cells and poorly differentiated Mahlavu cells. Cells were treated with CQ and the differentially expressed gene (DEG) data were acquired from a large multiplex panel of genes using the NanoString platform (Fig. 2d). Our experimental data indicated significant alterations in JAK/STAT, PI3K, RAS, MAPK pathways involving cytokine production in liver cells (full pathway list: Table S.3). These pathways were also presented in both KGs along with additional cytokine related pathways, such as interleukin signaling, along with dense connections to other biological components in COVID-19 CROssBAR-KGs, which is an expected output considering the mode of action of CQ in COVID-19 (Fig. 2c). Additional use-cases are provided in the Supplementary Information section 3. We believe that the CROssBAR system can be utilized towards the systematic analysis of the pharmacological effects of drugs as it brings relevant pieces of biological data together, that is relevant to the user’s query, which can be manually explored by the expert to build new hypotheses.

## 2. Methods (Online)

### 2.1. CROssBAR Database & API

#### Integrated data resources and the CROssBAR-DB

CROssBAR-DB has developed its bespoke ETL pipelines in Java 8 using the Spring batch framework to structure the jobs. The latter are executed on state-of-the-art EMBL-EBI LSF clusters powered by IBM in a parallel distributed fashion to reduce the processing time. The data are finally stored in MongoDB in the form of independent data collections, thus, providing schemaless flexibility and faster development, while sustaining data relationships in the form of nested documents. The pipelines have been both unit and integration tested using Spock framework in Groovy language.

The public databases integrated in the CROssBAR system can be listed along with the type of the biomedical data it contains as follows;

i. UniProt Knowledgebase (protein sequence and annotations including functions, domains, families, interactions, disease relations, pathway memberships, and more),
ii. IntAct (protein-protein interactions, currently incorporated directly from UniProt),
iii. InterPro (protein domain and family information),
iv. DrugBank (approved and investigational drugs and their targets),
v. ChEMBL (small molecule compounds, targets, bioassays and bioactivities collected from literature and other sources),
vi. PubChem (small molecule compounds, targets, bioassays and bioactivities collected from various resources),
vii. Reactome (pathway entries and their relations to proteins, currently incorporated directly from UniProt),
viii. KEGG (pathway and disease entries together with their relations to genes),
ix. Experimental Factor Ontology - EFO (disease terms integrated from multiple disease-centric databases including OMIM and Orphanet, and organized under an ontological system), and
x. Human Phenotype Ontology - HPO (phenotypic abnormality terms that relate to both genes and disease entries).

The statistics regarding the number of terms and annotations incorporated to CROssBAR-DB from each resource listed above is given in Fig. 1b. CROssBAR-DB schema is provided both in Supplementary Information Fig. S1 and in the GitHub repository of the project (https://github.com/cansyl/CROssBAR), where the attributes/fields belonging to each collection can be observed in full detail. KEGG is currently kept apart from the CROssBAR-API due to potential issues related to giving public access to the bulk of their data.

#### Obtaining biomedical data via the CROssBAR-API

CROssBAR data services (CROssBAR-API) are developed in Java 8 using Spring Boot’s web module in a RESTful architecture style. The API currently provides 12 endpoints documented using Swagger API documentation which allows endpoints to be tested within the documentation, and gives all information about the expected response schema. The API leverages the CROssBAR-DB hosted in a MongoDB platform to fetch data and filter results for users. The web services have been both unit and integration tested using Spock framework in Groovy language. CROssBAR-API is publicly available at https://www.ebi.ac.uk/Tools/crossbar/swagger-ui.html, where users can query independent collections over the indexed attributes/fields. The screenshots of the Swagger API are given in the Supplementary Information Fig. S3, where the 12 CROssBAR-DB collections are shown. Here, the collections entitled “Activities”, “Assays”, “Molecules” and “Targets” correspond to bioactivity, bioassay, compound and target entries in the ChEMBL database, respectively. “Drugs” corresponds to drug entries in DrugBank, “EFO disease terms” corresponds to the disease entries in the Experimental Factor Ontology, “HPO” corresponds to phenotype entries in the Human Phenotype Ontology, “Proteins” correspond to a subset of the protein entries in the UniProtKB. The remaining four collections belong to the PubChem data. There is no one-to-one correspondence between the incorporated data resources and the CROssBAR-DB collections since some of the resources had to be split to multiple collections for easier query (e.g., ChEMBL and PubChem). Also, some of the sources are directly incorporated from the UniProt database, thus, reside in the proteins collection (e.g., both terms and annotations for InterPro, Reactome, and only annotations for IntAct, OMIM and Orphanet).

It is possible to obtain cross-collection relational data (i.e., integrated relational data from multiple collections) by writing programmatic queries and submitting them to the API, as it is applied in CROssBAR-WS to construct the knowledge graphs. However, it is not possible to obtain this complex relational data in a single query using the Swagger graphical user interface. Currently, CROssBAR knowledge graphs do not include PubChem data due to both elevated computational demand (the sizes of PubChem collections are large) and high redundancy (a large portion of bioactivity data points in PubChem and ChEMBL databases are shared). However, it is possible to query the CROssBAR-DB using the provided API service, to obtain data entries from PubChem database collections. The database and API construction work has been handled by the Protein Function Development (UniProt database) team at EMBL-EBI, utilizing their expertise in biological database development and maintenance together with the available strong computational infrastructure, the team managed to build a huge but stable resource. The professional service providing approach applied by the team allowed the proper and constant maintenance of both the database and the entire CROssBAR system.

### 2.2. Deep-learning-based predictors and dataset construction

The identification of novel drug-like compounds and discovering new usages of existing drugs are key steps in drug discovery and development. Traditionally, this is accomplished via costly and time-consuming procedures and the rate of identifying novel drugs has decreased in recent years. Out of all different biomedical entity relation types, drug/compound-target protein interaction (DTI) is one of those with the highest rate of data incompleteness considering the current knowledge. There are more than 100 million distinct drug candidate compound records in total in public bioactive chemical databases such as ChEMBL and PubChem, let alone the theoretical number of all possible small molecules around 10^60^. Considering their pairwise combinations against hundreds of thousands of target biomolecules such as single proteins and macromolecular complexes, the current knowledge corresponds to less than 0.001% of the whole compound-target space^19^. The high rate of missing DTI data negatively impacts the integrated biomedical resources as well. In the CROssBAR project, we aimed to address this issue by producing machine learning based DTI predictions and incorporating these predictions to the CROssBAR resource. The studies specifically about the development of these tools have already been published or under review^11,12^, however, we used our tools to produce DTI predictions to be incorporated in our knowledge graphs in the framework of this study.

#### Bioactivity dataset construction

One critical topic in developing DTI prediction models is the source dataset to be used in system training procedures. It is especially critical to construct large-scale DTI datasets to train deep-learning models. To address this issue, we prepared a DTI dataset from the ChEMBL database that is suitable for training machine learning systems, with standardized filtering operations on targets, compounds and bioactivities. The dataset is periodically updated with each ChEMBL database release. We employed this dataset for the training and validation of the deep-learning based DTI prediction models we developed in the framework of the CROssBAR project, and also as the source dataset for drug/compound-target interaction space visualization (the methods are described below). It can also be used for developing new DTI prediction models. The current version of the bioactivity dataset (ChEMBL v27) is available for public use in: https://github.com/cansyl/CROssBAR/blob/master/CROssBAR_DB_API/ChEMBL27_preprocessed_activities_sp_b_pchembl.zip. Details regarding the dataset can be found in our recent article^11^.

#### Deep learning base predictor 1 - DEEPScreen

DEEPScreen was the first DTI prediction system that we developed in this endeavour. DEEPScreen is a high-performance drug–target interaction predictor that utilizes deep convolutional neural networks and 2-D structural compound representations (i.e., simple images) to predict their activity against intended target proteins. DEEPScreen system is composed of 704 target protein specific prediction models, each independently trained using experimental bioactivity measurements against many drug candidate small molecules, and optimized according to the binding properties of the target proteins. The main novelty of DEEPScreen is employing readily available 2-D structural representations of compounds at the input level instead of conventional drug/compounds descriptors (e.g., molecular fingerprints) that display limited performance. DEEPScreen produces binary predictions, meaning that a compound is either predicted as active or inactive against a target protein. During the development of this method, we also carried out cell-based *in vitro* wet-lab experiments on computationally generated DTI predictions, with the purposes of both validating the accuracy of the prediction models, and for gaining biological insight in the framework of health and disease, especially to contribute to the understanding of processes active in different cancer subtypes. DEEPScreen can be used for the fast screening of the chemogenomic space, to provide completely new DTIs that can later be investigated experimentally in the fields of drug discovery and repurposing^11^. The source code, datasets and the results of DEEPScreen are available at https://github.com/cansyl/deepscreen.

To enrich the DTI data in CROssBAR, DEEPScreen was employed to scan a considerable portion of the chemogenomic space and predicted more than 21 million new DTIs between 1.3 million drug candidate compounds in the ChEMBL database and 532 target proteins. A filtered version of these predictions (∼8 million) were incorporated in CROssBAR and displayed to users as part of CROssBAR-KGs. These predictions can directly be downloaded from: https://github.com/cansyl/CROssBAR/blob/master/CROssBAR_DB_API/CROssBAR_DEEPScreen_Largescale_DTI_predictions_filtered.tsv.zip.

#### Deep learning base predictor 2 - MDeePred

Our second deep-learning based DTI prediction system “MDeePred” adopts the proteochemometric approach, where both the compound and target protein features are employed at the input level to model their interaction, which enables the prediction of binders to under-studied or completely non-targeted proteins. In MDeePred, multiple types of protein features such as sequence, structural, evolutionary and physicochemical properties are incorporated within multi-channel 2-D vectors, which is then fed to state-of-the-art pairwise input hybrid deep neural networks, together with molecular fingerprint-based vectors of compounds. MDeePred predicts real-valued drug/compound-target protein interactions, which can be interpreted in terms of comparable response values such as IC50/Kd/Ki/potency^12^. The source code and datasets of MDeePred are available at https://github.com/cansyl/MDeePred.

In the framework of this study, we trained two MDeePred prediction models, with the aim of incorporating their DTI predictions to th COVID-19 CROssBAR-KG. One of these models was trained using ChEMBL experimental bioactivity data of orthologous ACE/ACE2 receptors from different organisms (i.e., human, rat, mouse and rabbit) and used for predicting new inhibitor drugs for human ACE2 receptor. The second model was trained using ChEMBL bioactivity data points that belong to 3C-like proteinase sub-unit of replicase polyprotein 1ab of closely related coronavirus strains (i.e., SARS, MERS, Feline and NL63 coronaviruses) and used for predicting new inhibitor drugs for SARS-CoV-2 3C-like proteinase. For both models, only ∼10,000 drug entries in the DrugBank database (the ones with investigational and approved drug status) were used as the query/test set, since the principle requirement for new potential COVID-19 treatments is to be exempt from early drug development procedures (e.g., pre-clinical analyses, phase I clinical trials, …). Five drugs with high predicted affinities (i.e., most of them with predicted IC50 < 2 uM for 3C-like proteinase and IC50 < 100 nM for ACE2) were selected for human ACE2 (i.e., 7-Hydroxystaurosporine, Eribaxaban, Becatecarin, Ticagrelor and Amcinonide) and for SARS-CoV-2 3C-like proteinase (i.e., Diloxanide furoate, Quinfamide, Phenyl aminosalicylate, Netarsudil and Amlodipine) and included in the COVID-19 CROssBAR-KG. Both the ChEMBL derived training datasets of these models and the full prediction results are provided in the GitHub repository of the CROssBAR project (https://github.com/cansyl/CROssBAR). The chemogenomic modelling approach used in MDeePred enabled us to provide predictions for these two targets, which would otherwise be impossible due to the unavailability of training data points, as SARS-CoV-2 3C-like proteinase has no experimental bioassay results in the source databases, and human ACE2 protein have only an insufficient number of experimental bioactivity data points.

### 2.3. Construction of knowledge graphs

In CROssBAR, the data is stored in a non-relational database (MongoDB), as separate collections for easy maintenance and fast querying. As a result, the database itself is not a knowledge graph. Instead, biologically relevant small-scale knowledge graphs are constructed on-the-fly, triggered by users’ queries with a single or multiple term(s) such as the names or ids of genes/proteins, diseases/phenotypes, compounds/drugs and/or pathways/biological processes of interest.

In CROssBAR knowledge graphs, biological entities are represented as vertices/nodes. Distinct types of nodes are defined for: *(i)* biomolecules (i.e., genes/proteins), *(ii)* biological mechanisms (i.e., processes/pathways), *(iii)* pathologies (i.e., diseases and phenotypes), and *(iv)* small molecule ligands used for treatment (i.e., drugs and drug candidate compounds). Relations between different types of biological entities are expressed by the edges of the graph. Edge types vary according to the defined relations. For a relation between; *(i)* two proteins, the edge is labelled as “interacts_with”, *(ii)* for a gene/protein and a disease, the edge label is “related_to”, *(iii)* for a drug/compound and a protein, the edge label is “targets”, *(iv)* for a gene/protein and a pathway, the edge label is “involved_in”, *(v)* for a gene/protein and a phenotype term, the edge label is “associated_with”, *(vi)* for a drug and a disease, the edge label is “indicates”, *(vii)* for a disease and a pathway, the edge label is “modulates”, and *(viii)* for a disease and a phenotype term, the edge label is “associated_with”.

The incorporation of pathway-related information (both signalling and metabolic pathways) in CROssBAR is done based on a membership-based approach, where pathways are expressed on the graph as single nodes and the nodes of those member proteins are connected to them via edges. This approach leaves out the detailed reaction-based mechanistic information provided in pathway databases such as Reactome and KEGG pathways; however, the inclusion of this information via applying a pathway resource styled network approach would prevent the generation of large heterogeneous networks composed of tens of different pathways and other components. Nevertheless, it is possible to explore these pathways in detail using the provided links, which takes the user to the corresponding page on that pathway database. Both Reactome and KEGG pathways provide the same type of biological information at the level of large-scale biological processes; however, Reactome also divides these processes into sub-pathways, whereas KEGG only provides the pathway information at a generic level. In CROssBAR, due to the way the overrepresentation analysis is done, only specific sub-pathways are incorporated from the Reactome database, in most cases. As a result, pathway information in the knowledge graphs is displayed at different levels of specificity, and thus, not redundant.

A simplified form of the knowledge graph construction work-flow is displayed in Fig. 1c. In this figure, the parts where the disease and gene/protein collections are queried are shown in full detail and queries on the rest of the components are simplified. The full-scale version of the knowledge graph construction procedure is displayed in Supplementary Information Fig. S2. Here, the finalized filtered dataset of each biological component (i.e., genes/proteins, diseases, phenotypes, drugs, compounds and pathways) is shown with a shape surrounded by a black frame. The graph is built using the entities in these datasets, together with their inter-component relations.

#### Node filtering via overrepresentation analysis

Due to the fact that the construction of the graph is based on including all biological components/terms that are connected to the query term(s) directly or indirectly in the database, without further filtering operations, most of the searches resulted in a huge graph composed of tens of thousands of nodes and hundreds of thousands of edges, including nearly each and every biological data entry in the source database. This kind of a graph would be unusable due to multiple reasons. First of all, it would not be possible to visually perceive a biologically relevant result from the giant network. Second, constructing and interactively displaying this graph would have computational requirements so high that it would not be feasible. To address this problem, we applied a multi-staged overrepresentation-based enrichment analysis process during the construction of graphs. In this analysis, we calculate an independent enrichment score for each biological entity in the database (i.e., a disease, phenotype, drug, compound, gene/protein or pathway), to be considered as its relevance to the graph that is being constructed. The calculation of enrichment score and its statistical significance is done using a modified version of the hypergeometric test for overrepresentation^20^, which also corresponds to a one-tailed Fisher’s exact test, and it is based on the statistics of the relations/connections with gene/protein nodes. For example, the enrichment score (*E*_*D,W*_) and its significance (*S*_*D,W*_), in terms of p-value, for a disease term *D*, for graph *W* is calculated as follows:

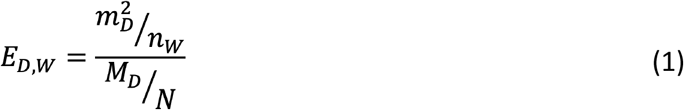

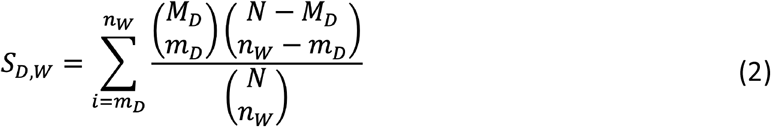

where *E*_*D,W*_ is the enrichment score calculated for the disease term *D* for graph *W; m*_*D*_^*2*^ represent the square of the number of genes/proteins in graph *W* that are associated with disease *D; n*_*W*_ represents the total number of gene/protein nodes in graph *W*; *M*_*D*_ is the total number of genes/proteins (not necessarily in graph *W*) that is associated with disease *D*; and *N* represents the total number of reviewed human gene/protein entries (i.e., UniProtKB/Swiss-Prot entries) in the CROssBAR database that is annotated with any disease entry. *S*_*D,W*_ represents the significance (p-value) for the disease term *D* for graph *W* calculated in the hypergeometric test.

While constructing the graph, an enrichment score is calculated for each disease entry in the CROssBAR database and these scores are used to rank disease entries according to their biological relevance to graph *W*. A cut-off value *k* is employed to include the top *k* relevant disease entries to graph *W*. The default value for *k* is 10, which means that only top-10 relevant diseases will be included in the graph. Apart from diseases, the same methodology is used to filter out the nodes of neighbouring genes/proteins, phenotypes, drugs, compounds and pathways. Significance values are not directly used in the filtering operation, since the main objective here is not only including significantly over-represented terms, but just reducing the number of nodes in the graph by filtering out the ones that are least relevant. In the traditional way of calculating an enrichment score, *m*_*D*_ is without square. The reason behind taking the square of *m*_*D*_ here is to break the tie between the scores of terms in favour of the one with a higher *m*_*D*_ value.

#### Formalizing the graph construction around gene/protein entries

During the construction of a knowledge graph, first, the gene/protein entries that are directly connected to the query term (i.e., core proteins) are fetched, such as the member genes/proteins of the queried signalling pathway. After that, neighbouring/interacting genes/proteins are added to the graph by calculating enrichment scores for each interacting protein, using the equation above, and filtering out based on the selected cut-off value. This is followed by the enrichment-based filtering and addition of other entity types; however, this time, both core and neighbouring genes/proteins are taken into consideration together to calculate the enrichment scores. If the user starts a heterogenous search that contains multiple terms from different entity types, both core and neighbouring genes/proteins are independently collected for each non-protein query term, queried gene/protein entries are added to this list (if there is any), and the entity collection process is continued using the union of these genes/proteins (Fig. 1c). This approach enables the exploration of direct and indirect relations between the queried terms.

#### Gene/protein filtering based on source organisms

We set a taxonomic filter for the inclusion of gene/protein entries in knowledge graphs, where the default selection is human (tax id: 9606), since the main focus of CROssBAR is biomedicine. Even though there are entries for proteins from hundreds of different organisms in the UniProtKB/Swiss-Prot database, only a few of these non-human protein entries possess annotations in terms of pathway memberships, targeting drugs/compounds and phenotype/disease implications. Thus, many of these protein entries are useless in terms of constructing biomedical knowledge graphs. Nevertheless, it is possible use the taxonomic filter in the web-service to include genes/proteins from a few additional organisms namely, Rattus norvegicus (Rat) [10116], Mus musculus (Mouse) [10090], Sus scrofa (Pig) [9823], Bos taurus (Bovine) [9913], Oryctolagus cuniculus (Rabbit) [9986], Saccharomyces cerevisiae (strain ATCC 204508 / S288c) (Baker’s yeast) [559292], Mycobacterium tuberculosis (strain ATCC 25618 / H37Rv) (MYCTU) [83332] and Escherichia coli (strain K12) (ECOLI) [83333].

#### Bioactive compound and bioactivity selection procedure

Small molecule compounds are selected and incorporated to KGs based on their reported bioactivities against target proteins. In a KG, a compound is represented as a node and a bioactivity is represented as an edge between a compound node and a gene/protein node. We start the bioactive compound collection procedure with a set of target gene/protein entries at hand (gathered in a previous step of the KG construction process), and obtain the compounds that are reported to be bio-actively interacting with these proteins, as their target biomolecules. Despite having a simple logic, this procedure is extremely complex due to practical reasons. Since there are more than 15 million bioactivity measurements in the ChEMBL database (v26), we rigorously filter these data points with the aim of providing only the most relevant bioactivity/compound information in CROssBAR-KGs. Since CROssBAR is a gene/protein centric system, we first filter out the data points where the target is not a single protein. We also set an organism filter for the targets, where the default selection is human. Additionally, we filter out bioactivities if their standard (activity) type is not one of these: IC50, EC50, AC50, XC50, Ki, Kd, potency; since these standard types provide roughly comparable measures of half-maximal response. Furthermore, we eliminated data points without a pChEMBL value, which standardizes the above-mentioned standard types under one value in the negative logarithmic scale. Bioactivity data points with an assigned pChEMBL value have usually received additional curation, and thus, they are more reliable. Finally, with the aim of only taking the data points at the active binding range (i.e., high affinities between the ligand and the target) we discard the data points with a pChEMBL value less than 5 (i.e., XC50 > 10 µM). Despite these filtering operations, we still usually end up with tens of thousands of compounds before the compound enrichment analysis, which significantly increases the KG construction run time. Exploiting the fact that it is a better choice to include a compound with higher binding affinity compared to a compound with a lower binding affinity for the same target protein, we set the pChEMBL value cut off value to 8, at the beginning of the compound collection procedure. Then, we reduce the cut off value and re-run the query if the total number of gathered compound entries is less than 1000 in the first run. We iteratively do this procedure until we obtain at least 1000 compound entries. Similarly, if the number of returned compounds is more than 2500 in the first run, we further increase the cut off iteratively until we obtain less than 2500 compound entries. This number (i.e., 1000 to 2500) is still much higher than the number of compounds we incorporate to a KG, which is between 0 and 50; however, we aim to enter the enrichment analysis with a high amount to be able to select the compounds that are interacting with multiple proteins in the network, not just one. Another reason is to be able to select diverse compounds, in terms of their scaffolds/structures, which is explained below (under the compound clustering sub-section).

#### Compound clustering

There are more than one million compound entries in the ChEMBL database, most of which have bioactivity data points against target biomolecules. Since it is not feasible to include each and every bioactive compound node in a KG (otherwise the graph would be extremely crowded), only the most overrepresented compounds are tried to be incorporated. We observed that some of the compounds with the same (or a very similar) enrichment score are also structurally very similar to each other. These are mostly molecules with matching scaffolds, which are screened against the same target and produced similar results in the same bioassay. Since their enrichment scores are similar as well, they are either selected or discarded together. To provide a better selection of compounds in the graph, we incorporated a structural property-based filtering in the enrichment analysis. The aim here is to select overrepresented compounds that are as diverse from each other as possible in terms of molecular structures, so that users will be provided with a variety of ligands for the target proteins in the graph. To achieve this, we calculated the pairwise molecular similarities between all compounds in CROssBAR-DB using circular fingerprints (ECFP4) and the Tanimoto coefficient. After that, we clustered the compounds based on a predefined similarity cut-off value of 0.5, meaning that each cluster is composed of compounds that are at least 50% similar to each other. The cluster information is pre-calculated and recorded on our server. Each time a knowledge graph is being constructed, enrichment score ranked compounds are checked one by one in terms of their cluster membership and if there already is a compound from the same cluster in the graph, the compound in turn is discarded (i.e., not incorporated to the graph). The same clustering-based selection approach is applied to incorporate compounds that are computationally predicted to interact with the proteins in the graph.

Following the finalization of the compound nodes, we check whether some of these compounds correspond to drugs that are already incorporated to the KG (since ChEMBL also contains bioactivity measurements belonging to approved or investigational drugs), using the identifier mapping between ChEMBL and DrugBank databases. When a positive case is detected, we merge these two nodes and set the node type as a drug, since drugs are considered more reliable in terms of evidence on their molecular properties and interactions, compared to drug candidate compound entries. If there are interactions reported both in DrugBank (as DTIs) and ChEMBL (as bioactivity measurements), we place all of the necessary edges from this drug node to the corresponding target protein nodes in the KG.

#### Evidence-based labelling of compound-target relations

Relations that signify the interactions between drugs/compounds and target proteins have three different sources with varying degrees of confidence. The most reliable relation in CROssBAR-DB is obtained from the DrugBank database, where the reported drug-target interaction (DTI) is verified by extensive analyses as part of an official drug development process. The resources that provide the interaction information that came second in terms of reliability are bioactivity databases such as ChEMBL. In these resources, reported bioactivities are obtained by experimental bioassays; however, they are not as extensively verified as in drug discovery and development procedures. The third in the list of interaction sources is the deep-learning based in-house computational predictions that we produced. These predictions are not verified by any experimental means so they should be considered with caution, even though we carried out numerous computational validation experiments for all predictions and provided *in vitro* experimental verification for a small selected set^11^. As a result, predicted relations comprise the least reliable part of the DTI information provided in CROssBAR-KGs. With the aim of transmitting this evidence-based relation confidence information to users, we used edge labels in the KGs. In terms of visualization, these labels are encoded on the graphs as colours, such that green colour corresponds to DTIs obtained from approved or investigational drugs, blue colour corresponds to experimental bioactivity measures obtained from ChEMBL, and the red colour corresponds to computationally predicted interactions. During the generation of KGs, if a specific relation is obtained from multiple sources (e.g., when the same relation is reported both in DrugBank and in ChEMBL) the edge label of the more reliable relation is incorporated. To accomplish this task, KG construction process comprises an edge label update procedure.

Another process we applied at this step is the edge addition. Some drugs possess bioactivity data points in ChEMBL in addition to their approved targets in DrugBank. To detect this, we first do a mapping between ChEMBL compound entries and DrugBank drug entries, to find the equivalent ChEMBL entry for each drug. After that, we identify the reported ChEMBL bioactivities between that compound and all of the proteins presented in the KG. For those relations, which were not already incorporated into the KG via DrugBank, we added blue coloured edges. The same procedure is applied for adding red coloured edges to the drugs and compounds that have extra computationally predicted target interactions.

### 2.4. CROssBAR Web-Service, User Interface and Layout

CROssBAR Web-Service (CROssBAR-WS) comprises both the backend and the frontend processes to construct the KGs and to display them to users. CROssBAR-WS comprises an underlying complex API query set that gathers data from the CROssBAR-DB. The underlying API query that collects relevant terms (entries) from 8 independent CROssBAR database collections together with their relations, is given in the GitHub repository of the project (https://github.com/cansyl/CROssBAR). CROssBAR-WS also contains a sophisticated graphical user interface that runs on our server. The open access technologies used in the construction of CROssBAR-WS are PHP, JavaScript (cytoscape.js), jQuery, CSS (BootStrap), MySQL.

A user query initiates the entity (node) gathering procedure first from the related database collections, using the CROssBAR RESTful API. Together with the entities that match the search term, the information regarding the related/connected entities are obtained from the corresponding collection. After that, the next database collection is queried with the terms gathered at the previous step. The order of the API queries follows the logic defined for the construction of the KGs, as given in Fig. 1c (simplified version), and in Fig. S2. (full version). Following the initiation of a query, the growing knowledge graph is displayed on the web-browser in real time (using CytoScape Web), starting from the collection and filtering of core and neighbouring genes/proteins. The process is continued with the collection, filtering and addition of phenotype/pathway/disease terms, drugs, bioactive compounds and predicted interacting compounds to the KG (as nodes), together with their relations with gene/protein nodes (as edges). The construction process is finished with the addition of respective edges between non-protein nodes.

An important subject in graph/network visualization is the layout. In CROssBAR-WS, we incorporated the standard layouts of CytoScape Web, such as circle, cose, grid and concentric. However, none of these layouts were sufficient for communicating highly heterogeneous graphs with 7 different types of nodes and 9 different types of edges. To address this problem, we developed the CROssBAR layout, in which biological terms (nodes) from a specific biomedical component (e.g., diseases, pathways, …) are placed on circular points within a fixed radius. With the aim of preventing overlapping nodes, the radius of each circle is selected as a different value. Curved edge style (i.e., unbundled-bezier) is applied to reduce the amount of edge crossing. More information regarding the usage of CROssBAR-WS and its user interface can be found at https://crossbar.kansil.org/tutorial.php.

CROssBAR web-service queries run in linear time and the actual duration of the process is correlated with the total number of core genes/proteins obtained within the query, together with the annotation volume of these genes/proteins. Highly studied genes/proteins usually have high number of associations, which in turn, extends the actual query runtime in practice. According to our tests, most of the queries with disease, gene/protein, drug and compound terms (in terms of both single and combinatory term searches) take between 1 and 3 minutes to complete (from job submission to the display of the whole graph). However, most pathway and some phenotypic term queries take longer, especially when the number of directly associated genes/proteins is over one hundred. With the aim of creating a better user experience, we applied a procedure in which the collected nodes and their edges are instantaneously and interactively displayed on the screen, before the end of the job. This way, users do not have to wait for the whole job to be finished before starting to explore the KGs.

The entire CROssBAR web-service including the web-site and the underlying API queries can be found in the GitHub repository of the project (https://github.com/cansyl/CROssBAR).

### 2.5. Generation of COVID-19 KGs

We constructed two versions of COVID-19 knowledge graph. First, the large-scale version that includes nearly the whole of the COVID-19 related information recently accumulated in the scientific literature, organized and presented in an interpretable way. Second, the simplified version, which is suitable for quick exploration. The aim behind constructing the simplified version was that the large-scale KG is not easily explorable visually due to its huge size. Since most of the COVID-19 related data has still not been integrated into the regular releases of biological databases, the data could not be pulled to the CROssBAR database at the time of writing this manuscript. As a result, we had to make manual interventions to obtain the data from CROssBAR data resources. We applied the same knowledge graph construction methodology incorporated in CROssBAR; however, we also conducted manual curation to a certain extent, since, unlike the rest of the CROssBAR data, COVID-19 related information has not been extensively curated yet. We saved the pre-constructed KGs, which are directly accessible and viewable through the links given on our web-service (https://crossbar.kansil.org/covid_main.php). It is also important to note that, due to the integrated data resources, CROssBAR heavily contains rare and complex disease data, and mostly leaves infectious diseases out. Nevertheless, the constructed COVID-19 graphs provided rich biomedical information. Below, we describe the methodology followed for the generation of CROssBAR COVID-19 KGs.

Construction of the large-scale COVID-19 graph started with acquiring the related EFO disease term named: “COVID-19” (id: MONDO:0100096). We also incorporated the disease term for “Severe acute respiratory syndrome” (id: EFO:0000694) (the original SARS) into the graph since SARS is better annotated compared to COVID-19. The full-scale COVID-19 KG construction is continued as follows:

#### COVID-19 related genes/proteins and PPIs

We obtained related genes/proteins and their interactions from the IntAct database’s COVID-19 dataset. Unlike a genetic disease, human genes/proteins represent only a portion of an infectious disease. Due to this, we aimed to incorporate SARS-CoV and SARS-CoV-2 genes/proteins, as well as, the host genes/proteins into the graph. Without any filtering, the KG contained 1,172 gene/protein and metabolite nodes from various organisms and 2,214 edges. Due to the high number of genes/proteins in the KG, there was a risk of incorporating non-specific/irrelevant terms from the other biological components at later steps. To address this risk, we applied several filtering operations on this data. First, we eliminated all non-gene/protein nodes and we discarded the genes/proteins if the corresponding organism is not human or SARS-CoV/SARS-CoV-2. Second, we eliminated the protein entries that are not reviewed (i.e., not from UniProtKB/Swiss-Prot) except SARS-CoV-2 ORF10 (accession: A0A663DJA2), which currently is an unreviewed protein entry in UniProtKB/TrEMBL. We also filtered out a portion of the host genes/proteins using interaction-based information, according to their confidence scores reported in IntAct. We discarded the edges between host proteins and SARS-CoV and/or SARS-CoV-2 proteins if the confidence score was less than 0.35. We also discarded the edges between host proteins in the KG (i.e., neighbouring proteins) if their interaction confidence score is less than 0.6. We removed the disconnected components made up of host proteins, which were formed due to the edge filtering operation. Orthology relations between SARS-CoV and SARS-CoV-2 genes/proteins were annotated with “is ortholog of” edge type. The subunits of large protein complexes such as the NSPs of replicase polyprotein 1ab of SARS-CoV/SARS-CoV-2 were mapped to their corresponding protein complex nodes with “is subunit of” edge label. After these operations, the finalized number of genes/proteins/subunits is 539 (475 host genes/proteins, and 33 SARS-CoV and 31 SARS-CoV-2 genes/proteins/subunits) and the number of edges (i.e., PPIs) is 1284. After this point, we started collecting new nodes and edges from various biological components based on the overrepresentation analysis and curation.

#### COVID-19 related drugs and compounds

The approved/investigational drug interactions of the COVID-19 related genes/proteins were retrieved from DrugBank database, v5.1.6 release. To incorporate only the most relevant drug-target interactions, a drug overrepresentation analysis was applied in terms of the target genes/proteins in the KG using the hypergeometric distribution, as described in the section entitled “Construction of knowledge graphs”. The selected drugs were mapped to the related protein targets in the graph using the edge label of green colour, as this represents the highest level of confidence in terms of ligand interactions. DrugBank also has a COVID-19 specific drug list, which includes a curated list of drugs currently under research for COVID-19 treatment. These drugs were included in the KG as well. Considering those without known targets (or the targets are known but not presented in the KG), we included them by connecting directly to the COVID-19 disease node. We also incorporated drug repurposing based curated and experimental results from new and critical SARS-CoV-2 related publications such as Gordon *et al*.^21^, and we mapped these interactions to our KG with suitable edge labels based on the data source. Finally, we added drug-disease relationships based on reported drug indications obtained from the KEGG resource. The KG contains well-studied drugs for COVID-19 treatment such as Chloroquine (DB00608), Remdesivir (DB14761), Favipiravir (DB12466), Dexamethasone (DB01234) and etc., as well as rather under-studied or non-studied ones (in the context of COVID-19) such as Isosorbide (DB09401) and Quercetin (DB04216).

For the retrieval of compound-target interactions based on experimentally measured bioactivities, ChEMBL database (v27) database was utilized. We retrieved the ChEMBL bioactivity data points in binding assays, where the targets are human or SARS proteins, and the pChEMBL value is greater than or equal to 5. Overrepresentation analysis was applied to select the most relevant ones. Here, only drugs/compounds with enrichment scores greater than 1 and p-value less than 0.05 were considered. Compounds were clustered based on Tanimoto coefficient based molecular similarities with a threshold of 0.5, and top 5 overrepresented compound nodes, that are in different clusters, were selected for each target protein and incorporated to the KG. We also incorporated selected compound - host target protein interactions from ChEMBL’s SARS-CoV-2 curated dataset. Finally, the edge labels are set accordingly (i.e., blue coloured edges).

For the computationally predicted drug and compound - target protein interactions, our in-house deep-learning-based tools DEEPScreen^11^ and MDeePred^12^ were used. DEEPScreen large-scale prediction run results were scanned and 326 bioactive drug/compound-target interaction predictions for 18 human proteins were incorporated to the KG following the application of overrepresentation analysis, similar to the one applied for selecting experimental bioactivities from ChEMBL. We trained two prediction models, one for human ACE2 receptor protein and one for SARS-CoV-2 3C-like proteinase using ChEMBL bioactivity datasets as our training dataset. Both models are used to scan full DrugBank drugs dataset to predict new binders for ACE2 and 3C-like proteinase to be utilized towards drug repurposing. The details of this process is given under the Methods sub-section entitled “Deep learning based predictors and dataset construction”. We only incorporated 5 selected inhibitors for each protein in the KG in order to avoid the crowding of the KG, however the full prediction sets are provided in the GitHub repository of the project (https://github.com/cansyl/CROssBAR). The selected bioactive drug predictions for ACE2 were Eribaxaban (DB06920), 7-Hydroxystaurosporine (DB01933), Becatecarin (DB06362), Ticagrelor (DB08816) and Amcinonide (DB00288); whereas the predictions for the 3C-like protease were Quinfamide (DB12780), Diloxanide furoate (DB14638), Phenyl aminosalicylate (DB06807), Netarsudil (DB13931) and Amlodipine (DB00381). The included predicted interactions were labelled with red coloured edges.

We also merged nodes with respect to drug-compound entry correspondences in DrugBank and ChEMBL databases. This way, some of the drug nodes also contain experimental bioassay-based relations (i.e., blue coloured edges) and computationally predicted relations (i.e., red coloured edges). At the end of these procedures, the total number of drugs (nodes) in the KG is 108 and the total number of drug interactions (edges) is 279. The total number of drug candidate small-molecule compounds in the KG is 233 and the total number of compound interactions (edges) is 517. Out of all drug/compound-target interaction edges, 135 correspond to drug development procedures, 335 to experimental bioassays and 326 to deep-learning-based predictions.

#### Pathways of COVID-19 related host genes/proteins

Signaling and metabolic pathway information was taken from Reactome (via CROssBAR database) and KEGG pathways data sources. The most relevant pathways were determined by the overrepresentation analysis and mapped to the related genes/proteins in the KG. Some of the incorporated pathways are directly related to SARS-CoV-2 infection such as “Viral mRNA Translation” (R-HSA-192823) or “ISG15 antiviral mechanism” (R-HSA-1169408) and innate pathways of the host such as “Endocytosis” (hsa04144), “Cell cycle” (hsa04110) or “NF-kappa B signaling pathway” (hsa04064). We also incorporated pathway-disease relations (in the sense of pathways that are modulated due to presence of certain diseases) based on the relation information obtained from the KEGG database. The finalized number of pathways in the KG is 57 (23 for KEGG and 34 for Reactome, among which there are corresponding terms) and the total number of gene/protein-pathway associations (edges) is 718 (244 for KEGG and 474 for Reactome).

#### COVID-19 related phenotypic implications

The resource for the phenotype terms is the Human Phenotype Ontology (HPO) database. For each phenotype term that is associated with at least one gene in the KG according to HPO data, we calculated an enrichment score and p-value via overrepresentation analysis. From the score-ranked HPO term list we selected phenotype terms that are not in a close parent-child relationship with each other in the HPO direct acyclic graph. HPO also has a curated list of SARS related phenotype terms. These terms were also added into the network and mapped to “COVID-19” and “Severe acute respiratory syndrome” disease nodes if their associated genes are not presented in the KG. This way, COVID-19 related phenotypes including symptoms such as Fever (HP:0001945), Myalgia (HP:0003326), Respiratory distress (HP:0002098), Immunodeficiency (HP:0002721) and etc. are included in the graph. The finalized number of phenotype terms in the KG (nodes) is 27 and the number of HPO term - gene/protein associations (edges) is 653. There are also 41 HPO term - disease associations.

#### Other associated diseases of COVID-19 related host genes/proteins

The aim behind this step is collecting the non-infectious (mostly genetic) diseases that utilize the same (or similar) biological mechanisms/processes of human, so that it may indicate potential risks for COVID-19 patients, or potential COVID-19 related repurposing options for drugs that are currently used to treat these associated diseases. For this, disease terms that are associated with genes/proteins in the COVID-19 KG are collected from the CROssBAR database resources: EFO disease collection (mainly including OMIM and Orphanet disease entries) and KEGG diseases database. The linkage of proteins and EFO terms was achieved through OMIM ids. The most relevant disease terms were selected based on the results of the overrepresentation analysis. Finally, disease-HPO term relations were also integrated into the KG using the disease association information provided in HPO resource. At the end of this step, diseases such as Combined oxidative phosphorylation deficiency (H00891), Amyotrophic lateral sclerosis - ALS (H00058), Bruck syndrome (H00514), Malignant Mesothelioma (EFO:1000355), and etc. have entered the KG. The finalized number of disease terms in the KG is 23 (10 for KEGG and 13 for EFO) and the number of disease - gene/protein associations (edges) is 52 (31 for KEGG and 21 for EFO).

The finalized large-scale COVID-19 KG includes 987 nodes (i.e., genes/proteins, drugs/compounds, pathways, diseases/phenotypes) and 3639 edges (i.e., various types of relations).

For the construction of the simplified COVID-19 KG, the starting point was the COVID-19 associated proteins in the UniProt COVID-19 portal (https://covid-19.uniprot.org/), instead of the IntAct SARS-CoV-2 interactions dataset, which was used as the base gene/protein set for the large-scale KG. The remaining steps of building the graph were mainly similar except that, additional nodes representing the organisms: human, SARS-CoV and SARS-CoV-2 were placed in the graph and connected to the corresponding proteins. The aim here was to prevent the presence of singleton protein nodes due to the reduced number of gene/protein nodes, and thus, the reduced number of PPIs, in the simplified graph. It is also important to note that the simplified version is not just a subset of the large-scale KG. Since the starting point of gene/protein collection were different in two KGs, the resulting graphs contain a slightly different content. For example, the drugs Siltuximab (DB09036), Pirfenidone (DB04951) and Lifitegrast (DB11611) are specific to the simplified KG. The simplified COVID-19 KG includes a total of 178 nodes and 298 edges. The detailed statistics for both KGs are provided in Table S.5.

For the Cytoscape network files of both COVID-19 KGs, overrepresentation analysis results, and for more information about the CROssBAR COVID-19 KGs please visit the CROssBAR project GitHub repository at: https://github.com/cansyl/CROssBAR. For directly visualizing and exploring the COVID-19 KGs interactively, please visit CROssBAR web-service at: https://crossbar.kansil.org/covid_main.php.

### 2.6. *In vitro* experimental procedures for Chloroquine treatment on liver cells

#### Cell Culture

Normal hepatocyte-like epithelial Huh7 cells and mesenchymal-like Mahlavu liver cells were grown in Dulbecco’s Modified Eagles Medium (DMEM) supplemented with 10% fetal bovine serum (FBS) (Gibco / Thermo Fisher Scientific), 0.1mM non-essential amino acids (Gibco / Thermo Fisher Scientific), and 100 Units/mL Penicillin/Streptomycin. Cells were maintained in 37°C in a humidified incubator under %5 CO_2_.

#### NCI-60 sulforhodamine B(SRB) cytotoxicity assay

Huh7 and Mahlavu liver cells were grown in 96-well plates (1000-200 cells/well) in an incubator for 24 hours. Both Mahlavu and Huh7 cells were treated with Chloroquine Phosphate (CQ) and water control in 40 μM to 0.3 μM concentrations for 72h. After fixation with cold 10% (w/v) trichloroacetic acid (MERCK) for an hour at +4°C, plate wells were washed 3 times with ddH_2_O. Each well was stained with 50μl of 0.4%SRB dye(Sigma-Aldrich) and incubated at RT for 10 min. To remove unbound SRB dye, wells were washed with 1% acetic acid for four times and left to air-dying. The protein-bound SRB was solubilized in 100 μl/well 10 mM Tris-Base solution, and the absorbance was measured with 96-well plate reader at 515 nm wavelength (ELx800, BioTek).

#### Gene expression analysis of Chloroquine with NanoString multiplex gene expression panel

Huh7 and Mahlavu liver cells were treated with CQ at cytotoxic doses of 3.6 μM and 12 μM, respectively, for 48 h. NanoString nCounter multiplex gene expression analysis, which includes 770 genes and various canonical pathways such as PI3K, MAPK, STAT, RAS, Cell cycle, DNA damage control, apoptosis, Hedgehog, Wnt, Transcriptional regulation, chromatin modification, and TGF-β, was applied on the RNA extracted from cells using the RNeasy mini kit (Qiagen), followed by the hybridization with code sets and scanning using the nCounter Digital Analyzer as instructed by the manufacturer (NanoString Technologies). Results were analysed using the Advanced Analysis Module on nSolver™3.0 software for quality control, normalization, and differential expression. The expression levels of each gene were normalized to those of control genes. After obtaining the differentially expressed gene list with the native software, a further filtering operation was applied based on the p-value (i.e., < 0.01), to identify the finalized list of genes with statistically significant expression changes. Differential expression of key pathways was revealed for both Huh7 and Mahlavu cell lines using the fold change of the member genes and their significance values. The resulting files are provided in the GitHub repository of the project (https://github.com/cansyl/CROssBAR).

## 4. Availability

CROssBAR web-service (including a tutorial on how to use the service) is available at: https://crossbar.kansil.org; CROssBAR database is available through our API at: https://www.ebi.ac.uk/Tools/crossbar/swagger-ui.html; all of the datasets, results and the source code of this project are available for download at: https://github.com/cansyl/CROssBAR; additional information is available in the CROssBAR project web-site at: https://cansyl.metu.edu.tr/crossbar.

## 5. Acknowledgements

This work was supported by the Newton/Katip Celebi Institutional Links program by TUBITAK, Turkey and British Council, UK (project no: 116E930, project acronym: CROssBAR), and the European Molecular Biology Laboratory core funds. We thank Dr Nurcan Tuncbag (faculty member, METU, Turkey), Dr Erden Banoglu (faculty member, Gazi University, Turkey), Dr Aybar Can Acar (faculty member, METU, Turkey) and Dr Tugba Suzek (faculty member, Mugla University, Turkey) for helpful discussions, comments and support. We also thank Dr Ian Dunham (Director, Open Targets, UK) and Dr Andrew Leach (Team leader, ChEMBL, EMBL-EBI, UK) for insightful discussions.

## 6. Author Contributions

T.D., R.C.A., M.M. and V.A., conceived the idea and planned the work. V.J., A.N., R.S., V.V., H.Z. and M.M constructed the database and the API. A.S.R., T.D. and V.A. developed the deep-learning-based prediction systems and produced large-scale relation predictions. H.A. and T.D. developed the knowledge graph construction methodology and prepared the datasets. A.A., T.D., R.C.A. and V.A. constructed the web-service. T.D., H.A., A.A. and A.S.R. performed the data analysis including the statistical tests. E.N. and R.C.A. carried out the *in vitro* experiments. T.D., H.A., V.J., R.S., R.C.A., M.M. and V.A. wrote and revised the manuscript. T.D., R.C.A., M.M. and V.A., supervised the overall study. All authors have approved the manuscript.

## 8. Additional information

Supplementary information is available for this paper.

