## Supplementary Information for "CROssBAR: Comprehensive Resource of Biomedical Relations with Deep Learning Applications and Knowledge Graph Representations"

#### 1. Background Work

There are numerous studies, tools and resources that integrate biological data (either from other data sources or by direct curation) and communicate it via textual or visual representations. One of the most commonly used resources in this sense are biological pathway databases such as Reactome[1], KEGG[2] and WikiPathways[3], where the interactions/reactions are communicated via network representations. STRING and STITCH databases are two well-known molecular interaction services, in which protein-protein and protein-chemical interactions are integrated from various resources, including both experimentally proven and electronically predicted data points, and presented to users as pre-computed networks[4, 5]. GeneMANIA is an online platform for exploring the relationships between genes over network representations generated by utilizing large-scale genomics and proteomics data, for gene prioritization and function prediction[6]. Apart from these well-known resources, there are other relevant studies in the literature. Relating the semantically same or similar terms from different ontological systems (without providing any visual output) was one of the earliest applications of the integration of structured biomedical data[7,8]. Another one of the early applications of heterogeneous biomedical data integration was the BioGraph data mining platform, which is shown to be successfully utilised for disease gene prioritization via random walks on the generated network[9]. In the Bio4j project, authors aimed to construct a graph-based platform as an infrastructure for integrating biological data. They stored public data obtained from sources such as UniProt, Gene Ontology and Expasy in independent graphs to be queried via domain specific languages such as Angulillos[10]. In project Rephetio, authors systematically integrated biomedical data from various resources and stored them in a graph database to construct the Hetionet resource, with the primary purpose of inferring new drug/compound–disease relations. Hetionet can be browsed by users via database queries in Cypher language[11]. With a similar approach, BioGrakn project aims to construct a biomedical knowledge graph using the Grakn database infrastructure. Users are required to download and run the system locally, which can be queried via the Grakl language[12]. Another system with the name, BioGraph, integrates gene/protein, function and cancer related miRNA data from various source databases and lets users to query the data using Gremlin query language to produce information on returned entities and simple network-based visualizations[13]. In a few studies, authors discussed alternative ways of extracting and relating biomedical data; *(i)* from the literature (i.e., articles and similar unstructured textual sources) via text mining[14-17], and *(ii)* from public biological databases providing ontological data via semantic integration[18], with the aim of constructing biomedical knowledge bases or graphs. Two recent studies evaluated and discussed the use of Wikidata, which is a community-driven semantic knowledge base, as data resource and infrastructure for the generation of biomedical knowledge graphs, over use cases and potential applications[19,20]. Lately, numerous bio-pharmaceutical companies start investing in biomedical data integration and mining using graph databases and knowledge graph-based representations[21,22].

Apart from the few widely adopted tools and services, many of the studies mentioned above, especially the ones dealing with heterogeneous biological data, suffer from issues that limit their functionality and/or usability. For example, some of them require highly specialized inputs from users, such as complex database queries, to generate the desired output, which may not be easy for researchers with little or no programmatic background. Some others do not provide an easily-interpretable visualization of the output data, which decreases their usability, since a complex corpus of information is usually hard to consume (without further computational analysis) when communicated via only textual or tabular definitions. Additionally, some resources are not properly maintained, causing the size and the content of the incorporated data to fall behind. In some cases, the authors just published a large dataset (e.g., a graph) for users to download, without any means of a relevant interactive sub-set generation, based on user queries. Finally, considering the proprietary resources belonging to pharmaceutical/ biodevelopment companies, which are claimed to fulfil most of the current biomedical data integration and representation requirements, these tools and services are only for internal utilization (in the course of their drug discovery and development projects), not open to public research.

Our literature review revealed that there is a critical requirement for fully open access, continuously updated and online biomedical data integration and knowledge graph representation tools/services with coding-free user interfaces, combined with cutting-edge artificial intelligence-based data enrichment applications, to be freely and easily used by the life-sciences research community.

#### 2. Investigation of COVID-19 Associated Drugs on the Graph

CROssBAR COVID-19 knowledge graph (KG) incorporates multiple drugs that can be utilized for developing novel treatments against SARS-CoV-2. Several of these drugs have already been reported in the COVID-19 literature and included based on this information; however, some of them were completely new. These new drugs have been incorporated to the graph either due to the overrepresentation analysis (based on the COVID-19 related host genes/proteins in the graph) or predicted to interact to with host or SARS-CoV-2 proteins by our deep-learning-based tools DEEPScreen and MDeePred. Here, we demonstrate a short literature-based validation study on the relevance of these new drugs for COVID-19. Table S.6. shows the promising drugs in our knowledge graph together with the source (i.e., whether they entered the graph due to enrichment or predicted by our deep-learning-based systems). It was interesting to observe that some of the drugs in this list are currently under clinical trials against COVID-19. The list includes calcineurin and IL-6 inhibitors such as cyclosporine, tocilizumab, amlodipine and siltuximab, which play roles in the immune system and effective against inflammatory diseases such as rheumatoid arthritis, juvenile idiopathic arthritis, and the Castleman disease. These disease entries are also incorporated into the KG as relevant diseases. A corticosteroid in the KG, prednisolone, is used to treat inflammatory conditions, and also associated with juvenile idiopathic arthritis. There are also other corticosteroids in the KG such as dexamethasone and methylprednisolone, that are associated with juvenile idiopathic arthritis. Since clinical studies indicate the efficacy of dexamethasone[23] and methylprednisolone[24,25] for the treatment of COVID-19, prednisolone could also be a promising option. Apart from these, other enriched drugs such as artenimol[26], lifitegrast[27], amcinonide[28], becatecarin[29] and quinfamide[30] have been shown as potential drugs for COVID-19 via docking studies, although these studies are yet to be peer-reviewed. Rocaglamide and its derivative didesmethylrocaglamide are involved in both full-scale and simplified COVID-19 KGs as enriched drugs. As a potent inhibitor of NF-κB activation in T-cells, rocaglamide and its derivatives may be also potential drug candidates for the treatment of COVID-19, though there is no COVID-19 related study about these drugs in the literature yet. It is also important to mention that, further research is required for assessing the potential of these drugs to be repurposed against SARS-CoV-2.

#### 3. Biomedical Data Exploration Use-Cases via the CROssBAR Web-Service

To provide an example about one of the many possible uses of the CROssBAR system, we explore the relation between a drug (trifluoperazine) and a disease (gastric cancer), to make a very quick and rough evaluation on the potential repurposing of this drug towards the disease of interest. Trifluoperazine is an approved antipsychotic agent mainly used in the treatment of schizophrenia. As far as we are aware, trifluoperazine has no *in vitro*, *in vivo* or clinical studies concerning the treatment of gastric cancer, although there are studies on other types of cancer such as colorectal[31], pancreatic[32], and lung[33], in the literature. Also, there is a study indicating the inverse association between antipsychotic use and the risk of gastric cancer[34]. Thus, this may be a convenient scenario for investigating the relationship between two potentially related biomedical entities, gastric cancer and trifluoperazine. To construct the corresponding knowledge graph, we queried the CROssBAR-WS with this drug and disease entries and selected the number of nodes to be incorporated to the graph (from each biomedical component) as 20. The resulting graph is shown in Fig. S4a.

Trifluoperazine exerts its antipsychotic effect with the blockage of dopamine D2 receptor. This relation is shown in the graph, where trifluoperazine binds to the DRD2 gene/protein node and is associated with the dopaminergic synapse pathway. In the KG, trifluoperazine also has other approved targets such as CALM1, ADRA1A and TNNC1 proteins (approved drug-target interaction edges have green colour), and these proteins are associated with calcium signalling pathway. Moreover, DRD2 and CALM1 are associated with the rap1 signalling pathway, as well. Both calcium and rap1 signalling pathways have other gene/protein associations such as ERBB2, KRAS, and CDH1, which are further associated with the gastric cancer disease. In the light of these relations, trifluoperazine can be explored via additional *in silico* and wet-lab studies, in terms of its potential to become a repurposed agent for the treatment of gastric cancer, which may show its activity on gastric cancer cells via calcium[35,36] and rap1 signalling pathways[37] (Fig. S4b).

Some of the proteins that are associated with the gastric cancer (e.g., KRAS, ERBB2, TP53, etc.) are also related to other cancer disease nodes in the graph such as the pancreatic cancer, ovarian cancer, endometrial cancer and cholangiocarcinoma, which means that trifluoperazine may also have a potential against these cancer types, worthy of further exploration. Other antipsychotic or anxiolytic agents such as risperidone, haloperidol, perphenazine, buspirone, droperidol, and prochlorperazine are enriched in the network as well, which bind to DRD2, CALM1 and/or ADRA1A. These drugs may also become alternative repurposed drugs for gastric cancer treatment or other cancers presented in the KG. In addition to the above-mentioned approved drug-target interactions, the graph also includes enriched drugs and drug like compounds having experimentally measured bioactivities - from ChEMBL- (shown with blue coloured edges) or computationally predicted interactions - by our in-house tool DEEPScreen- (shown with red coloured edges) against the targets DRD2, ADRA1A, EBP, and SIGMAR1; which can also be considered for the diseases in the graph. Finally, there are several phenotypic implication terms (from HPO) on the KG, such as the abnormal urine carbohydrate level and the congenital hypertrophy of retinal pigment epithelium, which are associated with gastric cancer disease node and/or gastric cancer related genes. These phenotypic implications could also be helpful for disease diagnosis.

Apart from the exploration of the potential drug repurposing applications, a drug search on CROssBAR can also be utilized towards identifying new drug-like compounds with similar target-based bioactivities. This kind of exploration can be useful for medicinal chemists and other researchers working on drug discovery. It is generally accepted that compounds with highly similar molecular structures also have similar bioactivities; however, there is no generally accepted approach for identifying compounds that can be alternatives to an approved drug, when there is no structural similarity between the drug and the candidate compounds. In this example, we query Sorafenib (https://www.drugbank.ca/drugs/DB00398) on CROssBAR-WS, which is a drug approved for the treatment of primary kidney and primary liver cancers, to construct the knowledge graph that includes a relevant set of biomedical data. An interesting observation on the resulting KG are the compound nodes: CHEMBL272938 (https://www.ebi.ac.uk/chembl/compound_report_card/CHEMBL272938/) and CHEMBL3910171 (https://www.ebi.ac.uk/chembl/compound_report_card/CHEMBL3910171/), which contain high number of shared targets with Sorafenib (5 out 10 of the approved target proteins of Sorafenib are also the targets of these compounds) indicated by the bioassay-based interactions (blue coloured edges on the graph) for CHEMBL272938 and by computationally predicted interactions (red coloured edges on the graph) for CHEMBL3910171. It is also important to note that, the other half of the approved targets of Sorafenib could also be shared with these compounds; however, we currently do not have further information about it. High overlap between these targets indicate the potential of these compounds to be an alternative for Sorafenib. It is also important to note that this result could not be obtained with a conventional molecular similarity search, as Sorafenib, CHEMBL272938 and CHEMBL3910171 have highly dissimilar structures (neither a substructure search, nor a pairwise molecular similarity search -with the minimum similarity threshold of 40%- on the ChEMBL database could not detect any similarity between these three molecules).

#### 4. CROssBAR's Alignment with FAIR Principles

At each step of developing the CROssBAR system, we considered the alignment with FAIR data principles[38] to contribute to the movement towards fully open access, standard and easily (re)usable biomedical data. Since we mainly integrate data from other open access biomedical resources, we did not create new identifiers; however, we extensively share the identifiers in the source databases together with links, with the aim of making our data findable. Considering accessibility, CROssBAR system is reachable through various ways including the all open access API and web-service (interactively displaying the knowledge graphs) and through the project repositories (e.g., https://github.com/cansyl/CROssBAR) which includes the source codes and all datasets. CROssBAR data is highly interoperable as it mainly integrates heterogeneous biomedical data that is normally scattered throughout many independent data resources, and presents it in a coherent and standardized form. CROssBAR is released under the Creative Commons Attribution 4.0 (CC BY 4.0) license which mostly prevents restrictions on the downstream reuse of data, thus aligning it with the reusability item of FAIR. It is also important to note that, since CROssBAR is mainly obtaining its data from other open access biomedical data services, their compliance with FAIR principles affects CROssBAR's alignment as well, even though this effect is only indirect. For this reason, we plan to encourage biomedical data providers to consider prioritizing FAIR compliance while developing their own resources.

#### 5. Conclusion

In this work, we presented CROssBAR, a new system that consumes large-scale biomedical data from various resources and distils it to present a coherent piece of information, relevant to a user-queried biomedical concept, communicated via heterogeneous knowledge graphs. To build CROssBAR, we constructed a NoSQL database and stored the data, developed deep-learning based models to enrich the data at hand by predicting missing links, conducted wet-lab experiments to evaluate the relevance of our computational analysis, built networks of integrated information containing genes/proteins, diseases/phenotypes, biological processes/pathways and drugs/compounds, and presented all these to the life-sciences research community in an open access, user friendly web-service. We hope that our well-maintained and continuously developed service will be well adopted by researchers from various domains to aid their investigations by exploring high-level relations across different biomedical concepts.

As future work, we plan to integrate additional biomedical resources to CROssBAR such as the cell-type/tissue based transcriptomics and proteomics data and other omics based (e.g., glycomics, lipidomics, etc.) annotations for biomolecules, genomic variation data especially to be associated with disease implications, evolutionary information such as paralogy and orthology relations across species, biomolecular function annotations (e.g., GO terms, EC numbers), data sources and tools for infectious diseases and their mechanisms, and literature-based information (e.g., articles) as evidences of incorporated relations. Furthermore, we plan to incorporate evidence-based edge weights in our knowledge graphs and also to the overrepresentation analysis procedure, with the aim of assessing the relevance of each biological term to the user's query, with higher specificity. We also plan to incorporate *in silico* relation predictions between different layers of the biomedical data, in addition to compound/drug-target protein interactions, such as gene/protein-disease/phenotype, gene/protein-function and drug-disease relations, by both constructing novel in-house prediction systems and by incorporating approaches from the literature, including the state-of-the-art graph-based link prediction. It is also important to state that, we continuously focus on providing a well-maintained, fast, scalable and practical service to the life-sciences research community with periodic back-end and front-end improvements.

26. Moussa Sehailia S. C. In-silico Studies of Antimalarial-agent Artemisinin and Derivatives Portray More Potent Binding to Lys353 and Lys31-Binding Hotspots of SARS-CoV-2 Spike Protein than Hydroxychloroquine: Potential Repurposing of Artenimol for COVID-19. Preprint at https://chemrxiv.org/articles/preprint/In-silico_Studies_of_Antimalarial-agent_Artemisinin_and_Derivatives_Portray_More_Potent_Binding_to_Lys353_and_Ly

s31-Binding_Hotspots_of_SARS-CoV-2_Spike_Protein_than_Hydroxychloroquine_Potential_Repurposing_of_Artenimol_for_COVID-19_/12098652/1 (2020).

40. Liao, Y., Wang, J., Jaehnig, E. J., Shi, Z., & Zhang, B. WebGestalt 2019: gene set analysis toolkit with revamped UIs and APIs. *Nucleic acids research, 47*(W1), W199-W205 (2019).

### 7. Supplementary Figures

#### Figure S.1

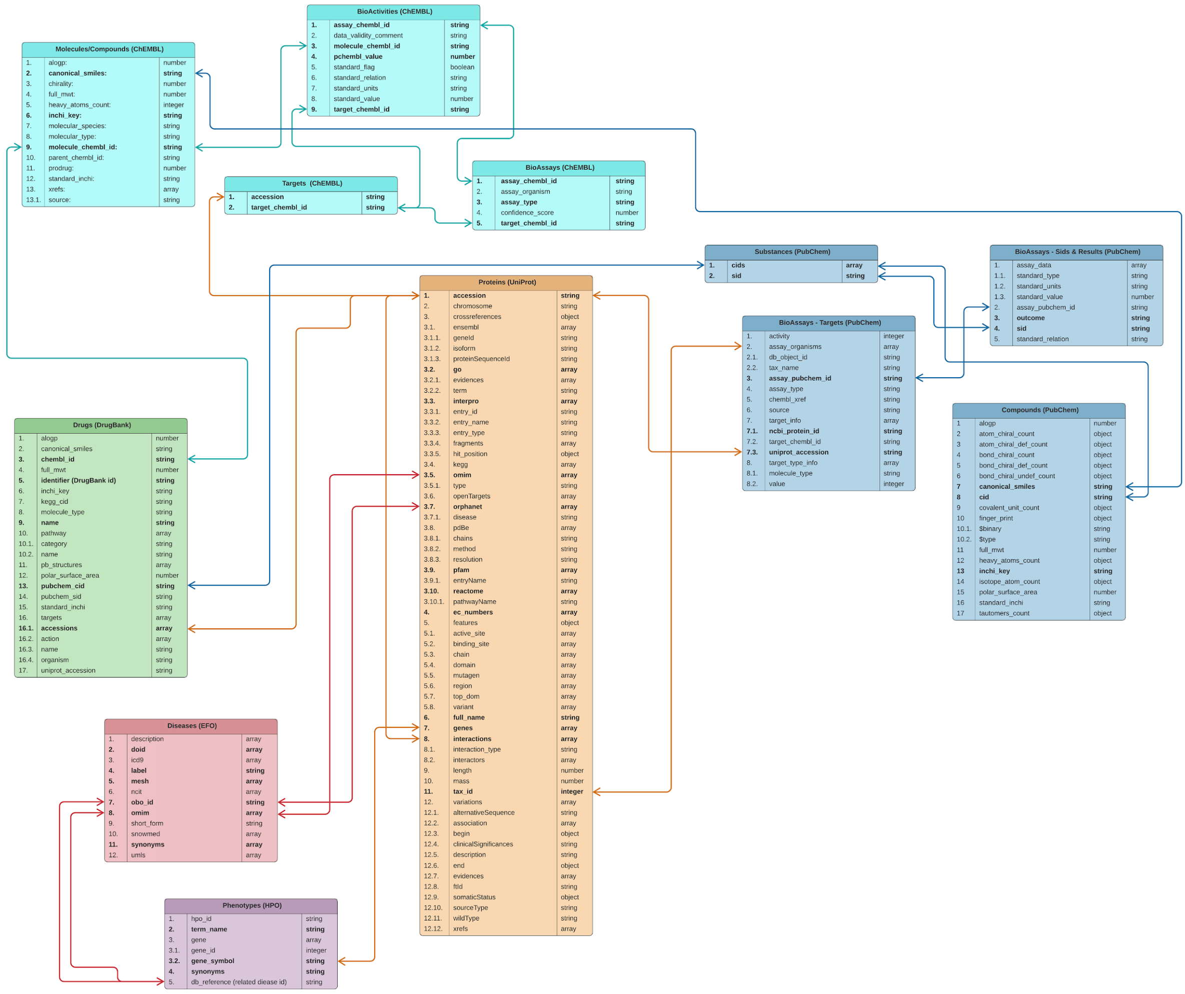


**Figure S.1.** Schematic representation of the independent collections of CROssBAR Mongo noSQL database, displaying the cross-collection relations.

#### Figure S.2

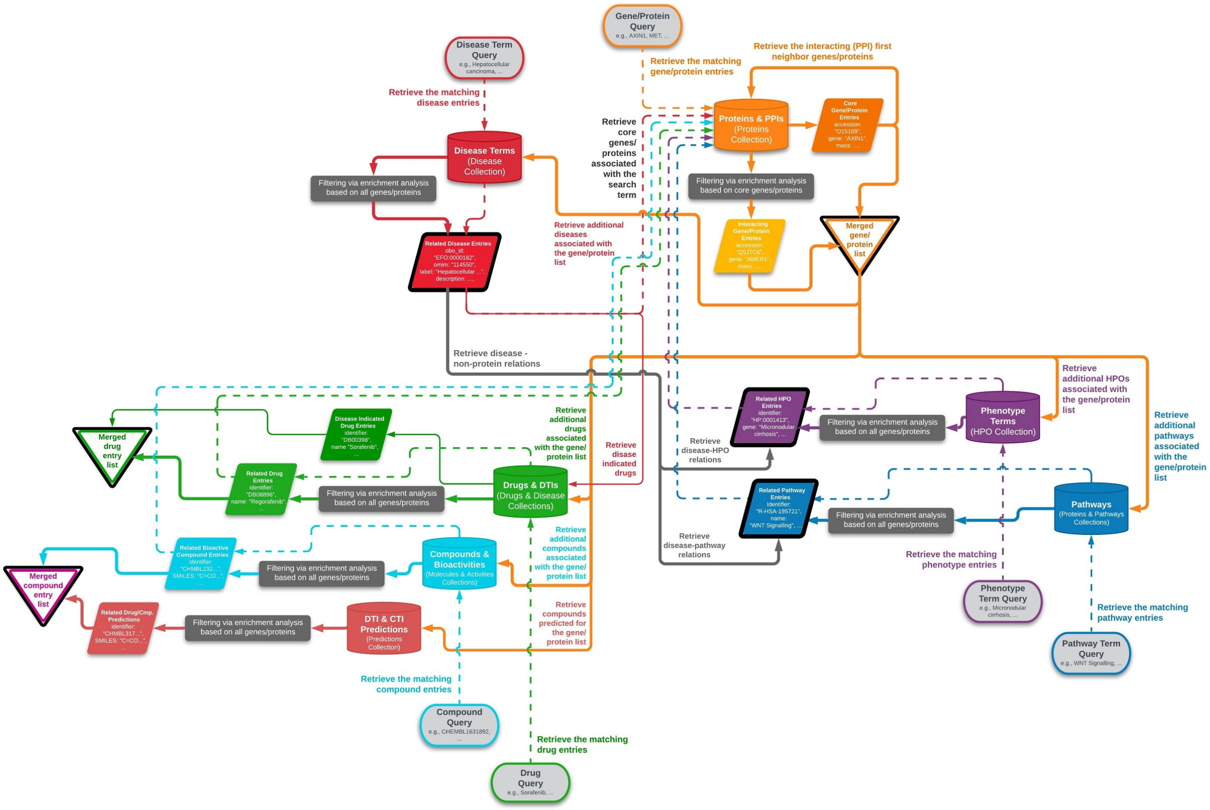


**Figure S.2.** Full-scale work-flow of the CROssBAR knowledge graph construction process.

#### Figure S.3

(a)

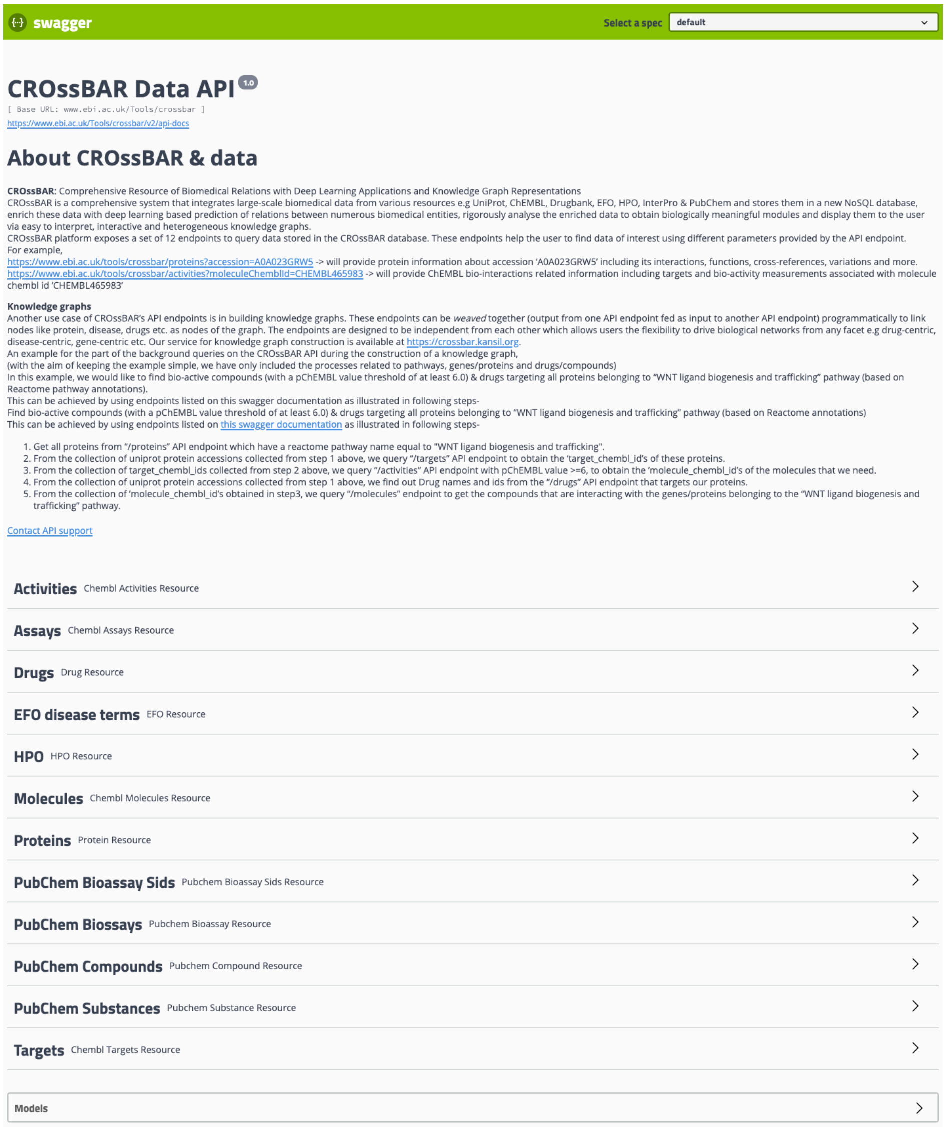

swagger
Select a spec default

### CROssBAR Data API 1.0

[ Base URL: www.ebi.ac.uk/Tools/crossbar ]
https://www.ebi.ac.uk/Tools/crossbar/v2/api-docs

#### About CROssBAR & data

**CROssBAR**: Comprehensive Resource of Biomedical Relations with Deep Learning Applications and Knowledge Graph Representations
CROssBAR is a comprehensive system that integrates large-scale biomedical data from various resources e.g UniProt, ChEMBL, Drugbank, EFO, HPO, InterPro & PubChem and stores them in a new NoSQL database, enrich these data with deep learning based prediction of relations between numerous biomedical entities, rigorously analyse the enriched data to obtain biologically meaningful modules and display them to the user via easy to interpret, interactive and heterogeneous knowledge graphs.
CROssBAR platform exposes a set of 12 endpoints to query data stored in the CROssBAR database. These endpoints help the user to find data of interest using different parameters provided by the API endpoint.
For example,
https://www.ebi.ac.uk/tools/crossbar/proteins?accession=A0A023GRW5 -> will provide protein information about accession 'A0A023GRW5' including its interactions, functions, cross-references, variations and more.
https://www.ebi.ac.uk/tools/crossbar/activities?moleculeChemblId=CHEMBL465983 -> will provide ChEMBL bio-interactions related information including targets and bio-activity measurements associated with molecule chembl id 'CHEMBL465983'

**Knowledge graphs**
Another use case of CROssBAR's API endpoints is in building knowledge graphs. These endpoints can be *weaved* together (output from one API endpoint fed as input to another API endpoint) programmatically to link nodes like protein, disease, drugs etc. as nodes of the graph. The endpoints are designed to be independent from each other which allows users the flexibility to drive biological networks from any facet e.g drug-centric, disease-centric, gene-centric etc. Our service for knowledge graph construction is available at https://crossbar.kansil.org.
An example for the part of the background queries on the CROssBAR API during the construction of a knowledge graph,
(with the aim of keeping the example simple, we have only included the processes related to pathways, genes/proteins and drugs/compounds)
In this example, we would like to find bio-active compounds (with a pChEMBL value threshold of at least 6.0) & drugs targeting all proteins belonging to "WNT ligand biogenesis and trafficking" pathway (based on Reactome pathway annotations).
This can be achieved by using endpoints listed on this swagger documentation as illustrated in following steps-
Find bio-active compounds (with a pChEMBL value threshold of at least 6.0) & drugs targeting all proteins belonging to "WNT ligand biogenesis and trafficking" pathway (based on Reactome annotations)
This can be achieved by using endpoints listed on this swagger documentation as illustrated in following steps-

1. Get all proteins from "/proteins" API endpoint which have a reactome pathway name equal to "WNT ligand biogenesis and trafficking".
2. From the collection of uniprot protein accessions collected from step 1 above, we query "/targets" API endpoint to obtain the 'target_chembl_id's of these proteins.
3. From the collection of target_chembl_ids collected from step 2 above, we query "/activities" API endpoint with pChEMBL value >=6, to obtain the 'molecule_chembl_id's of the molecules that we need.
4. From the collection of uniprot protein accessions collected from step 1 above, we find out Drug names and ids from the "/drugs" API endpoint that targets our proteins.
5. From the collection of 'molecule_chembl_id's obtained in step3, we query "/molecules" endpoint to get the compounds that are interacting with the genes/proteins belonging to the "WNT ligand biogenesis and trafficking" pathway.

Contact API support

**Activities** Chembl Activities Resource
**Assays** Chembl Assays Resource
**Drugs** Drug Resource
**EFO disease terms** EFO Resource
**HPO** HPO Resource
**Molecules** Chembl Molecules Resource
**Proteins** Protein Resource
**PubChem Bioassay Sids** Pubchem Bioassay Sids Resource
**PubChem Biossays** Pubchem Bioassay Resource
**PubChem Compounds** Pubchem Compound Resource
**PubChem Substances** Pubchem Substance Resource
**Targets** Chembl Targets Resource

**Models**

(b)

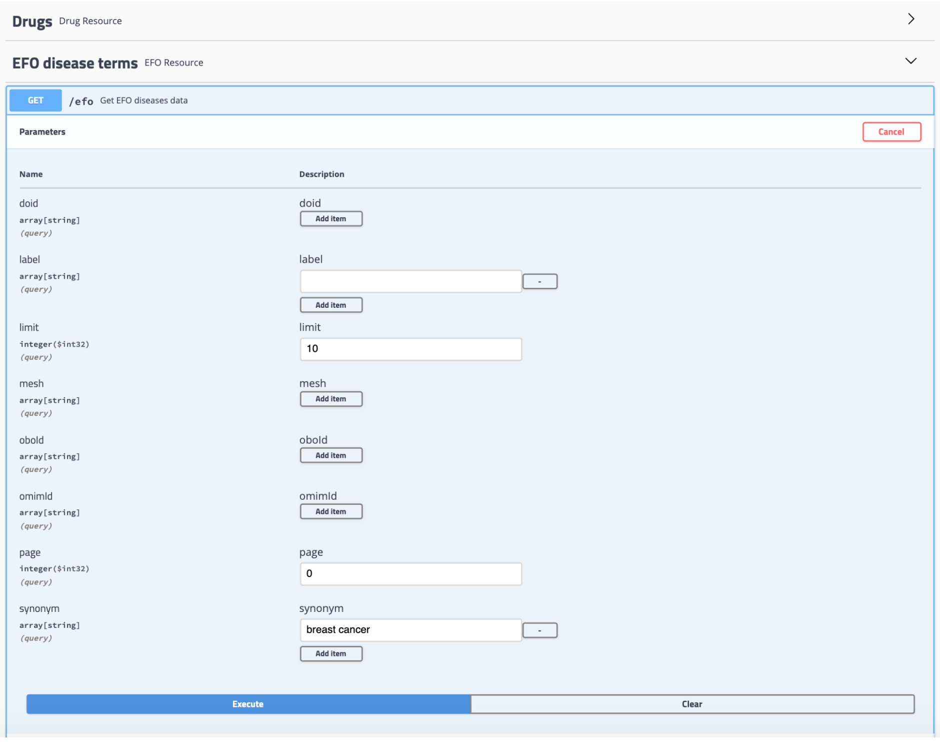

(c)

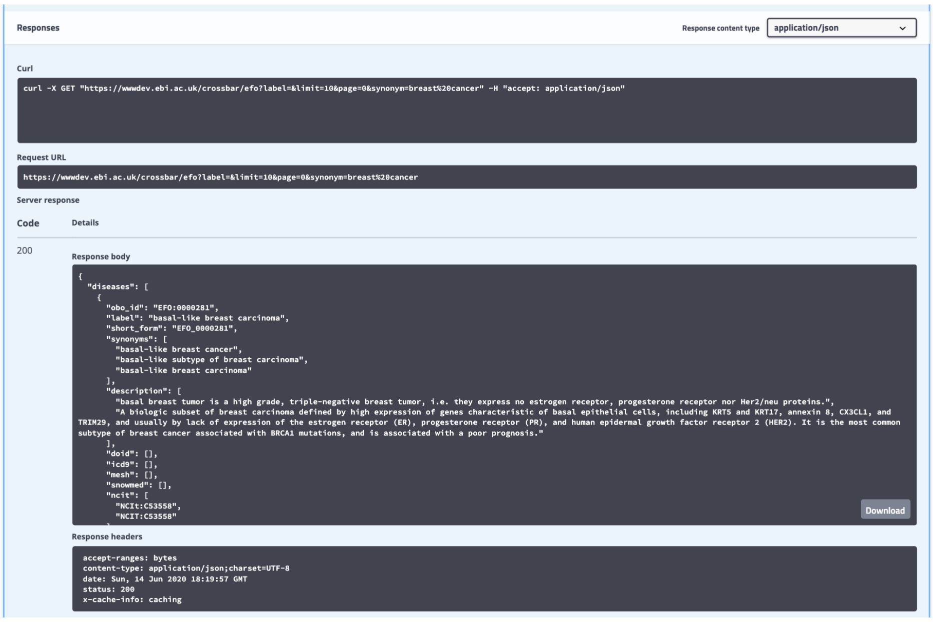

**Figure S.3.** CROssBAR API Swagger web interface (https://wwwdev.ebi.ac.uk/crossbar/swagger-ui.html); **(a)** names of 12 endpoints at the main page; **(b)** an example API query on EFO disease terms collection with "breast cancer" searched as the synonymous term name; **(c)** query response: the same query in command line and as URL, and the list of resulting database entries that match the query together with their attributes.

#### Figure S.4

**(a)**

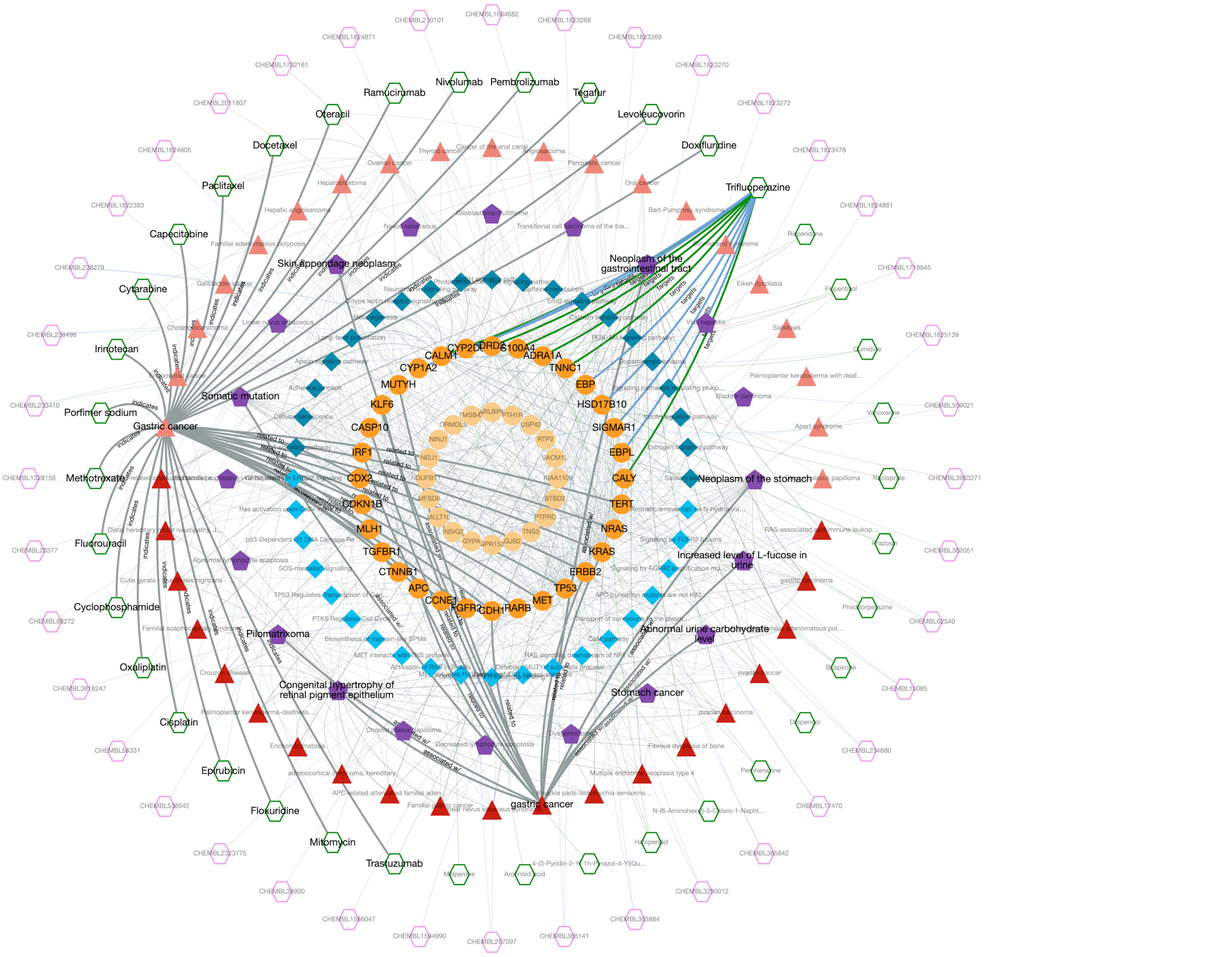
Nivolumab
Pembrolizumab
Ramucirumab
Tegafur
Oteracil
Levoleucovorin
Docetaxel
Doxifluridine
Paclitaxel
Trifluoperazine
Capecitabine
Skin appendage neoplasm
Neoplasm of the gastrointestinal tract
Cytarabine
Irinotecan
Somatic mutation
Porfimer sodium
Gastric cancer
Methotrexate
Neoplasm of the stomach
Fluorouracil
Increased level of L-fucose in urine
Cyclophosphamide
Abnormal urine carbohydrate level
Pilomatrixoma
Oxaliplatin
Congenital hypertrophy of retinal pigment epithelium
Stomach cancer
Cisplatin
Epirubicin
Floxuridine
gastric cancer
Mitomycin
Trastuzumab
CYP2D6
DRD2
S100A4
ADRA1A
CALM1
TNNC1
CYP1A2
MUTYH
EBP
HSD17B10
KLF6
SIGMAR1
CASP10
IRF1
EBPL
CALY
CDX2
CDKN1B
NRAS
TERT
MLH1
KRAS
TGFBR1
ERBB2
CTNNB1
TP53
APC
CCNE1
MET
FGFR2
CDH1
RARB

**(b)**

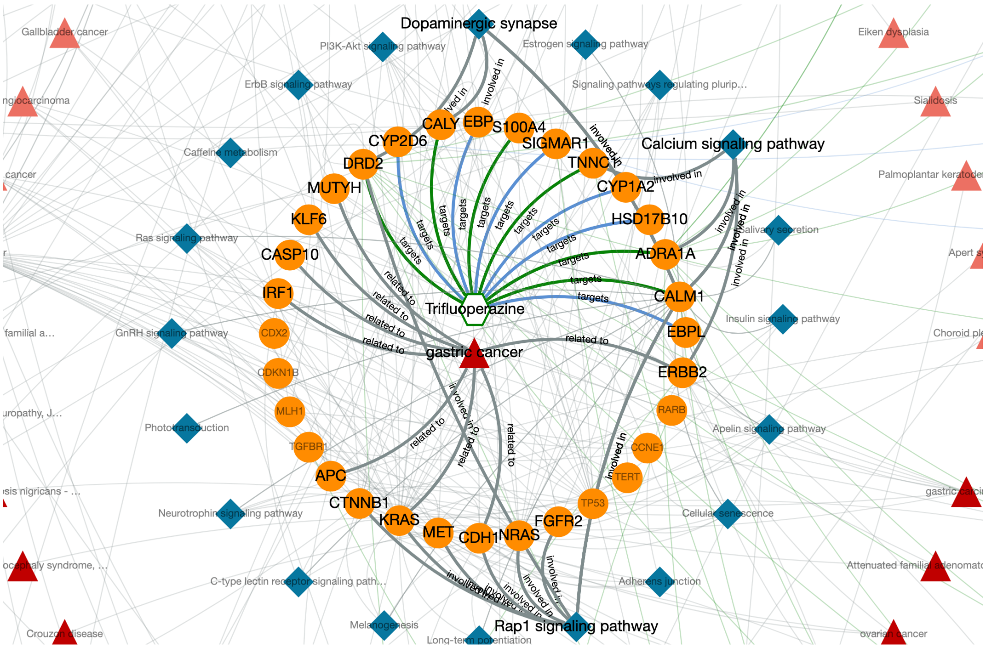
Dopaminergic synapse
Gallbladder cancer
PI3K-Akt signaling pathway
Estrogen signaling pathway
Eiken dysplasia
ErbB signaling pathway
Signaling pathways regulating plurip...
Sialidosis
CALY
EBP
S100A4
CYP2D6
SIGMAR1
Calcium signaling pathway
Caffeine metabolism
DRD2
TNNC1
CYP1A2
Palmoplantar keratode
MUTYH
HSD17B10
KLF6
Salivary secretion
Ras signaling pathway
CASP10
ADRA1A
Apert s
IRF1
Trifluoperazine
CALM1
Insulin signaling pathway
GnRH signaling pathway
CDX2
EBPL
Choroid pl
gastric cancer
CDKN1B
ERBB2
MLH1
RARB
Phototransduction
Apelin signaling pathway
TGFBR1
CCNE1
APC
TERT
CTNNB1
TP53
Neurotrophin signaling pathway
Cellular senescence
KRAS
MET
CDH1
NRAS
FGFR2
Attenuated familial adenomato
C-type lectin receptor signaling path...
Adherens junction
Melanogenesis
Rap1 signaling pathway
Crouzon disease
ovarian cancer
Long-term potentiation
targets
related to
involved in

**(c)**

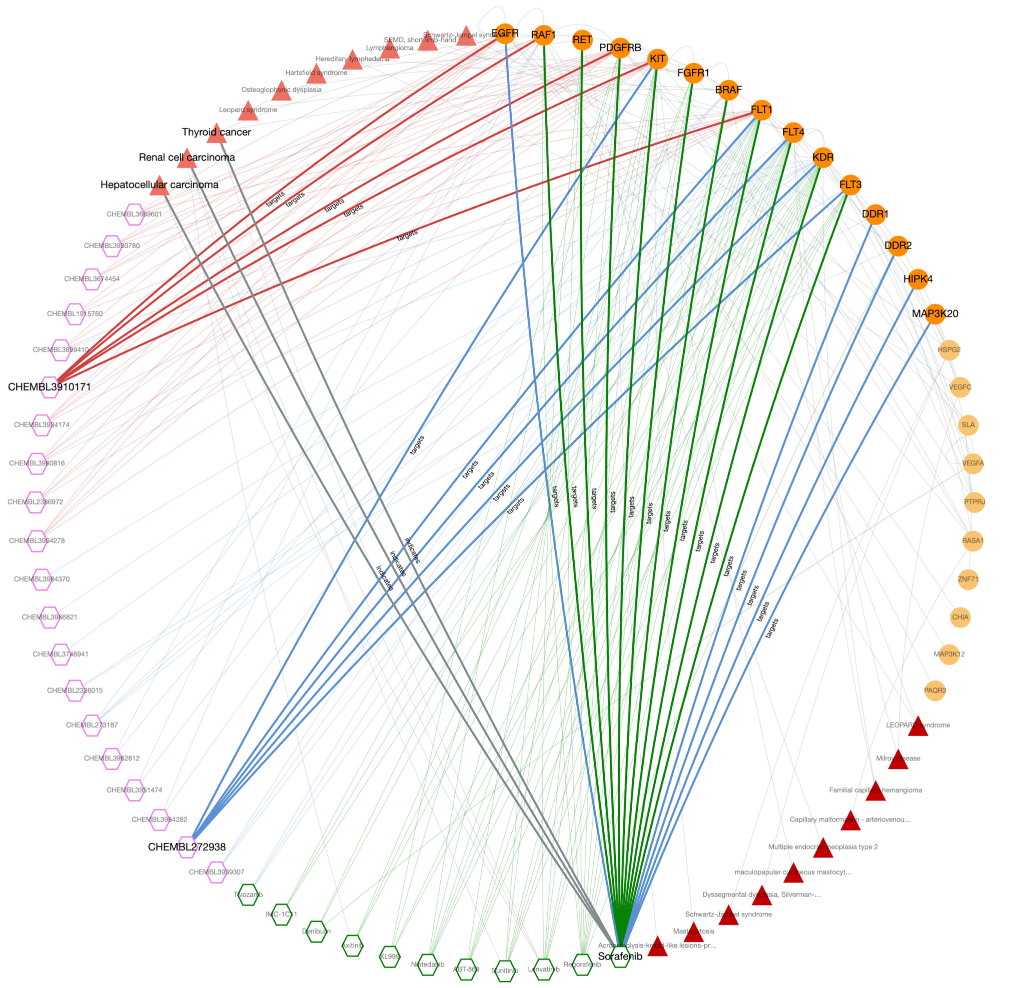


**Figure S.4.** Example cases of data exploration using the CROssBAR web-service, **(a)** the output knowledge graph of trifluoperazine and gastric cancer query; **(b)** critical signalling pathways and their relation to trifluoperazine and gastric cancer over critical genes/proteins; and **(c)** target interaction similarity between structurally dissimilar molecules: Sorafenib, CHEMBL272938 and CHEMBL3910171 (circular layout).

#### 8. Supplementary Tables

**Table S.1** CROssBAR database statistics.

| High-level data component group | Specific biomedical component | Number of entries |
|---|---|---|
| Genes/Proteins (UniProt) | Genes | 144,924 |
| | Proteins | 575,402 |
| | Proteins w/ PPI | 21,353 |
| | PPIs | 83,173 |
| | Domain Annotations | 948,635 |
| Bioactivities (ChEMBL) | Bioassays | 1,060,283 |
| | Compounds | 1,828,820 |
| | Bioactivities | 15,207,914 |
| | Targets | 12,091 |
| Bioactivities (PubChem) | Bioassays | 1,067,568 |
| | Compounds | 97,351,543 |
| | Substances | 252,877,567 |
| | Bioactivities | 268,562,338 |
| Drugs (DrugBank) | Drugs | 13,563 |
| | Targets | 4,914 |
| | DTIs | 19,272 |
| Diseases (KEGG+EFO) | EFO Disease Terms | 5,366 |
| | EFO Genes (Unique #) | 4,626 |
| | EFO Annotations | 13,851 |
| | KEGG Disease Terms | 1,901 |
| | KEGG Disease Genes (Unique #) | 3,710 |
| | KEGG Disease Annotations | 6,040 |
| Phenotypes (HPO) | HPO Terms | 9,061 |
| | HPO related Genes (Unique #) | 4,317 |
| | HPO Annotations | 580,607 |
| Pathways (KEGG+Reactome) | KEGG Pathway Terms | 337 |
| | KEGG Pathway Genes (Unique #) | 8,040 |
| | KEGG Pathway Annotations | 32,516 |
| | Reactome Terms | 7,762 |
| | Reactome Proteins (Unique #) | 68,898 |
| | Reactome Annotations | 208,213 |

**Table S.2.** List of the differentially expressed genes (DEGs) of chloroquine phosphate treated liver cells (Huh7 and Mahlavu), with a p-value cut-off < 0.01.

**Cell line: Huh7**

| Gene name | Log2 fold change | p-value | probe.ID |
|---|---|---|---|
| ABL1 | 0.183 | 0.000546 | NM_005157.3:3200 |
| ANGPT1 | -0.171 | 0.00108 | NM_001146.3:2080 |
| ARID1A | 0.203 | 0.000134 | NM_006015.4:5495 |
| ATM | -0.25 | 0.000887 | NM_138292.3:6688 |
| B2M | 0.132 | 0.00608 | NM_004048.2:25 |
| BAMBI | -0.171 | 0.00145 | NM_012342.2:832 |
| BAP1 | -0.149 | 0.00279 | NM_004656.2:240 |
| BCL2L1 | -0.192 | 1.66E-05 | NM_138578.1:1560 |
| BCOR | -0.369 | 0.00449 | NM_001123383.1:1630 |
| BID | 0.189 | 0.00203 | NM_197966.1:2095 |
| BNIP3 | 0.149 | 0.000208 | NM_004052.2:325 |
| BRCA2 | 0.0707 | 0.00607 | NM_000059.3:115 |
| C19orf40 | 0.355 | 3.51E-06 | NM_152266.3:376 |
| CAPN2 | -0.513 | 0.00282 | NM_001748.4:2085 |
| CASP8 | -0.105 | 0.00304 | NM_001228.4:301 |
| CCNA2 | -0.0804 | 0.00201 | NM_001237.2:1210 |
| CCNB1 | -0.0811 | 0.00251 | NM_031966.2:715 |
| CCNE2 | 0.218 | 0.00156 | NM_057735.1:50 |
| CDC25A | 0.0634 | 0.00529 | NM_001789.2:690 |
| CDC6 | -0.0589 | 0.00767 | NM_001254.3:1300 |
| CDH1 | 0.164 | 0.00136 | NM_004360.2:1230 |
| CDK2 | -0.179 | 0.00541 | NM_001798.2:220 |
| CDK4 | 0.104 | 0.00765 | NM_000075.2:1055 |
| CDKN2D | -0.554 | 0.0024 | NM_001800.3:870 |
| CEBPA | 0.137 | 0.00866 | NM_004364.2:1320 |
| COL2A1 | 0.417 | 1.56E-05 | NM_001844.4:4745 |
| CREB3L3 | 0.294 | 0.000332 | NM_001271995.1:1050 |
| CREB5 | -0.228 | 0.00118 | NM_182898.2:1885 |
| CSF3R | 0.675 | 0.00275 | NM_156038.2:90 |
| CUL1 | 0.111 | 1.11E-05 | NM_003592.2:1487 |
| DAXX | -0.117 | 0.0084 | NM_001350.3:1875 |
| DDIT3 | 0.5 | 1.98E-05 | NM_004083.4:40 |
| DKK1 | -0.16 | 0.000474 | NM_012242.2:75 |
| DNMT1 | -0.075 | 0.00034 | NM_001379.2:1495 |
| DUSP4 | 0.282 | 0.000107 | NM_057158.2:3115 |
| DUSP5 | 0.128 | 0.000351 | NM_004419.3:675 |
| E2F5 | -0.0998 | 0.00168 | NM_001951.3:444 |
| EFNA1 | 0.285 | 0.00906 | NM_004428.2:650 |
| EIF4EBP1 | -0.0695 | 0.0105 | NM_004095.3:363 |

| Gene name | Log2 fold change | p-value | probe.ID |
|---|---|---|---|
| IL6R | 0.558 | 0.00014 | NM_000565.2:993 |
| IL8 | 0.321 | 1.32E-05 | NM_000584.2:25 |
| ITGA2 | -0.115 | 0.00566 | NM_002203.2:475 |
| ITGB8 | 0.383 | 8.61E-05 | NM_002214.2:2609 |
| JAG1 | -0.145 | 0.00865 | NM_000214.2:915 |
| JAK2 | 0.222 | 0.0055 | NM_004972.2:455 |
| KAT2B | 0.229 | 0.00372 | NM_003884.3:1220 |
| KDM5C | 0.0422 | 0.000153 | NM_004187.2:1170 |
| KMT2C | 0.136 | 0.00696 | NM_170606.2:12505 |
| KMT2D | -0.127 | 0.000176 | NM_003482.3:6070 |
| LFNG | -0.755 | 0.000944 | NM_001040168.1:717 |
| LIFR | 0.0762 | 0.00956 | NM_002310.3:2995 |
| MAD2L2 | -0.223 | 6.49E-05 | NM_001127325.1:290 |
| MAML2 | -0.806 | 0.00055 | NM_032427.1:4125 |
| MAP2K1 | -0.0886 | 0.00458 | NM_002755.2:970 |
| MAP3K14 | 0.506 | 0.000158 | NM_003954.1:620 |
| MAPK8 | 0.0894 | 0.00051 | NM_002750.2:945 |
| MCM2 | -0.148 | 0.00264 | NM_004526.2:2945 |
| MNAT1 | 0.0794 | 0.00721 | NM_002431.2:975 |
| MSH6 | -0.0825 | 0.00904 | NM_000179.1:3525 |
| MYC | 0.0783 | 0.0014 | NM_002467.3:1610 |
| NASP | -0.221 | 0.000635 | NM_172164.1:2970 |
| NF1 | -0.0983 | 0.00692 | NM_000267.2:1035 |
| NFE2L2 | -0.178 | 0.000167 | NM_006164.3:995 |
| NFKBIZ | 0.308 | 0.000801 | NM_001005474.1:2030 |
| NOTCH1 | 0.0941 | 0.00144 | NM_017617.3:735 |
| NPM1 | -0.383 | 0.000825 | NM_002520.5:10 |
| NRAS | -0.0928 | 0.000532 | NM_002524.3:877 |
| NTHL1 | -0.241 | 0.00124 | NM_002528.5:476 |
| NUMBL | 0.364 | 0.00397 | NM_004756.3:591 |
| NUPR1 | 1.1 | 0.00249 | NM_001042483.1:829 |
| PAK7 | -0.28 | 0.00347 | NM_177990.1:615 |
| PDGFA | -0.558 | 0.00911 | NM_002607.5:2460 |
| PDGFC | -0.243 | 7.72E-06 | NM_016205.1:10 |
| PDGFRB | -0.198 | 0.00372 | NM_002609.3:840 |
| PIK3R3 | 0.216 | 0.000351 | NM_003629.3:1800 |
| PIM1 | 0.478 | 0.000563 | NM_002648.2:1630 |
| PLAU | 0.474 | 0.000477 | NM_002658.2:793 |
| POLD1 | -0.394 | 0.00403 | NM_002691.2:2392 |

| | | | |
|---|---|---|---|
| ENDOG | -0.106 | 0.00731 | NM_004435.2:694 |
| EP300 | 0.139 | 0.00345 | NM_001429.2:715 |
| EPHA2 | -0.208 | 0.00263 | NM_004431.2:1525 |
| EPOR | 0.314 | 0.000218 | NM_000121.2:1295 |
| ERBB2 | 0.127 | 0.00756 | NM_004448.2:2380 |
| ETV1 | -0.145 | 0.00276 | NM_004956.4:1719 |
| ETV4 | -0.153 | 0.0027 | NM_001079675.1:1535 |
| FANCB | 0.423 | 0.00158 | NM_152633.2:2470 |
| FANCE | -0.466 | 0.000715 | NM_021922.2:1275 |
| FANCG | -0.201 | 0.00355 | NM_004629.1:1900 |
| FGFR2 | 0.503 | 0.00163 | NM_000141.4:647 |
| FGFR3 | -0.0863 | 0.0065 | NM_022965.2:3170 |
| FLNA | -0.362 | 0.000317 | NM_001456.3:7335 |
| FN1 | -0.0797 | 0.000226 | NM_212482.1:1776 |
| FOS | 1.05 | 1.68E-05 | NM_005252.2:1475 |
| FST | -0.141 | 0.000331 | NM_006350.2:575 |
| FUT8 | 0.247 | 0.000113 | NM_004480.4:2841 |
| GADD45A | 0.345 | 7.75E-05 | NM_001924.2:865 |
| GNA11 | -0.164 | 0.00319 | NM_002067.1:555 |
| GNAQ | -0.165 | 0.00862 | NM_002072.2:1100 |
| GNAS | 0.0653 | 0.00911 | NM_080425.1:1910 |
| GNG12 | -0.151 | 0.000221 | NM_018841.3:245 |
| GNG4 | -0.279 | 0.00275 | NM_004485.2:215 |
| GNG7 | 0.24 | 0.00458 | NM_052847.1:3920 |
| GTF2H3 | 0.0392 | 0.00241 | NM_001516.3:70 |
| H3F3A | -0.108 | 0.000457 | NM_002107.3:190 |
| HDAC1 | -0.183 | 0.000104 | NM_004964.2:785 |
| HIST1H3H | -0.0973 | 7.93E-05 | NM_003536.2:355 |
| HMGA1 | -0.183 | 0.00252 | NM_145904.1:871 |
| HMGA2 | -0.111 | 0.00275 | NM_003484.1:328 |
| HOXA9 | 0.289 | 0.00278 | NM_152739.3:1015 |
| HSPA2 | -0.347 | 0.00214 | NM_021979.3:2095 |
| ID2 | 0.0544 | 0.00489 | NM_002166.4:505 |
| IDH1 | 0.2 | 0.00181 | NM_005896.2:105 |
| IGFBP3 | -0.351 | 0.000131 | NM_000598.4:1255 |
| IKBKG | 0.379 | 8.18E-05 | NM_003639.2:470 |
| IL1R1 | 0.362 | 0.000269 | NM_000877.2:4295 |

| | | | |
|---|---|---|---|
| POLR2H | -0.126 | 0.00098 | NM_001278698.1:940 |
| PPARG | -0.263 | 0.000212 | NM_015869.3:1035 |
| PPP3CA | -0.319 | 0.000434 | NM_000944.4:3920 |
| PRKCA | -0.123 | 0.000426 | NM_002737.2:5560 |
| PRKDC | -0.075 | 0.00354 | NM_006904.6:12750 |
| PRKX | -0.341 | 0.000311 | NM_005044.1:2590 |
| PRLR | -0.184 | 0.000673 | NM_001204318.1:563 |
| PROM1 | -0.238 | 0.00121 | NM_006017.1:925 |
| PTTG2 | -0.127 | 0.000152 | NM_006607.2:5 |
| RAD50 | 0.147 | 0.00924 | NM_005732.2:5397 |
| RAF1 | 0.0629 | 0.00789 | NM_002880.2:1990 |
| RELA | -0.16 | 0.00378 | NM_021975.3:1990 |
| RELN | 0.222 | 0.000153 | NM_005045.2:345 |
| RHOA | -0.212 | 0.00033 | NM_001664.2:1230 |
| RUNX1 | 0.416 | 0.0044 | NM_001754.4:635 |
| SETD2 | 0.188 | 0.000112 | NM_014159.6:6160 |
| SHC2 | 0.331 | 0.001 | NM_012435.2:698 |
| SMAD2 | 0.145 | 0.000271 | NM_001003652.1:4500 |
| SMAD4 | 0.0814 | 0.00388 | NM_005359.3:1370 |
| SMARCB1 | -0.118 | 0.0036 | NM_003073.3:1060 |
| SOCS1 | -0.389 | 2.06E-05 | NM_003745.1:1025 |
| SPRY1 | -0.434 | 0.000495 | NM_005841.1:810 |
| SPRY2 | -0.107 | 0.00232 | NM_005842.2:85 |
| STK11 | -0.2 | 0.00263 | NM_000455.4:2060 |
| TBL1XR1 | 0.0898 | 0.00555 | NM_024665.4:915 |
| TCF7L1 | 0.3 | 0.00317 | NM_031283.1:2215 |
| TFDP1 | -0.134 | 0.000719 | NM_007111.4:1826 |
| TGFB1 | -0.361 | 0.00153 | NM_000660.3:1260 |
| TGFBR2 | -0.121 | 0.000536 | NM_001024847.1:1760 |
| THBS4 | 0.25 | 0.0078 | NM_003248.3:985 |
| TNFAIP3 | 0.488 | 9.29E-05 | NM_006290.2:260 |
| TP53 | -0.154 | 0.000392 | NM_000546.2:1330 |
| TSC1 | 0.322 | 0.00115 | NM_000368.3:100 |
| UBE2T | 0.0624 | 0.00199 | NM_014176.3:595 |
| VEGFA | 0.266 | 0.000124 | NM_001025366.1:1325 |
| WIF1 | 0.883 | 3.42E-05 | NM_007191.2:765 |
| XRCC4 | 0.209 | 0.000844 | NM_003401.3:772 |

#### Cell line: Mahlavu

| Gene name | Log2 fold change | P-value | probe.ID |
|---|---|---|---|
| ABL1 | 0.196 | 0.000142 | NM_005157.3:3200 |
| ALKBH2 | 0.23 | 0.00364 | NM_001001655.2:907 |
| ALKBH3 | 0.0718 | 0.00275 | NM_139178.3:690 |
| ANGPT1 | 0.183 | 0.00135 | NM_001146.3:2080 |
| APC | -0.24 | 0.000634 | NM_000038.3:6850 |

| Gene name | Log2 fold change | P-value | probe.ID |
|---|---|---|---|
| LAMB3 | 0.343 | 0.00263 | NM_000228.2:695 |
| LEF1 | 0.118 | 0.00617 | NM_016269.3:1165 |
| LEPR | -0.397 | 0.000642 | NM_001003679.1:2000 |
| LIF | 0.779 | 0.00119 | NM_002309.3:1240 |
| LTBP1 | 0.593 | 0.000799 | NM_000627.3:4124 |

| | | | |
|---|---|---|---|
| ARID1A | -0.284 | 0.00124 | NM_006015.4:5495 |
| ARID2 | -0.0882 | 0.0102 | NM_152641.2:3355 |
| ASXL1 | -0.0805 | 0.000481 | NM_001164603.1:472 |
| ATR | -0.248 | 0.00643 | NM_001184.2:565 |
| B2M | 0.169 | 0.0046 | NM_004048.2:25 |
| BAP1 | -0.0531 | 0.00866 | NM_004656.2:240 |
| BCL2L1 | 0.0837 | 0.00122 | NM_138578.1:1560 |
| BNIP3 | 0.409 | 4.03E-06 | NM_004052.2:325 |
| BRAF | -0.123 | 0.00297 | NM_004333.3:565 |
| BRCA1 | 0.186 | 0.00578 | NM_007305.2:1275 |
| BRCA2 | 0.206 | 0.00445 | NM_000059.3:115 |
| BRIP1 | 0.43 | 0.000703 | NM_032043.1:1130 |
| CASP7 | 0.25 | 0.00341 | NM_001227.3:915 |
| CASP8 | 0.0783 | 0.0053 | NM_001228.4:301 |
| CBL | -0.211 | 0.000837 | NM_005188.2:7485 |
| CCNB1 | -0.132 | 0.000125 | NM_031966.2:715 |
| CCNB3 | -0.0904 | 0.000954 | NM_033671.1:35 |
| CCND1 | 0.452 | 6.21E-05 | NM_053056.2:690 |
| CDC25B | -0.184 | 7.60E-05 | NM_021873.2:3045 |
| CDC25C | -0.233 | 0.00309 | NM_001790.2:1055 |
| CDC6 | 0.233 | 0.000373 | NM_001254.3:1300 |
| CDK2 | 0.324 | 0.00183 | NM_001798.2:220 |
| CDK4 | 0.128 | 7.50E-05 | NM_000075.2:1055 |
| CDKN1A | 0.824 | 1.87E-05 | NM_000389.2:1975 |
| CDKN1B | -0.24 | 0.000518 | NM_004064.2:365 |
| CDKN2C | -0.264 | 0.00369 | NM_001262.2:1295 |
| CDKN2D | -0.234 | 0.000906 | NM_001800.3:870 |
| CHEK2 | 0.121 | 0.00526 | NM_007194.3:140 |
| COL5A1 | 0.0662 | 0.00984 | NM_000093.3:6345 |
| CREB3L1 | -0.113 | 0.000205 | NM_052854.1:195 |
| CTNNB1 | -0.0484 | 0.00303 | NM_001904.3:2265 |
| CYLD | 0.192 | 3.54E-05 | NM_015247.1:2890 |
| DDB2 | 0.362 | 0.000916 | NM_000107.1:840 |
| DDIT3 | 0.365 | 2.00E-05 | NM_004083.4:40 |
| DDIT4 | 0.564 | 0.000469 | NM_019058.2:85 |
| DUSP4 | 0.813 | 1.73E-05 | NM_057158.2:3115 |
| DUSP5 | 0.486 | 5.60E-06 | NM_004419.3:675 |
| DUSP6 | 0.836 | 0.000328 | NM_001946.2:1535 |
| E2F1 | -0.083 | 0.00364 | NM_005225.1:935 |
| EFNA5 | 0.395 | 0.000258 | NM_001962.2:5035 |
| EGFR | 0.152 | 0.000696 | NM_201282.1:360 |
| EIF4EBP1 | 0.172 | 8.06E-06 | NM_004095.3:363 |
| ENDOG | -0.137 | 5.00E-04 | NM_004435.2:694 |
| EP300 | -0.104 | 0.00266 | NM_001429.2:715 |

| | | | |
|---|---|---|---|
| MAD2L2 | -0.0859 | 0.00528 | NM_001127325.1:290 |
| MAML2 | 0.227 | 0.00547 | NM_032427.1:4125 |
| MAP2K1 | 0.216 | 0.00127 | NM_002755.2:970 |
| MAP2K4 | -0.247 | 0.000241 | NM_003010.2:2830 |
| MAP2K6 | -0.287 | 0.000167 | NM_002758.3:555 |
| MAP3K1 | -0.0658 | 0.00628 | NM_005921.1:2525 |
| MAPK1 | -0.0858 | 0.00232 | NM_138957.2:430 |
| MAPK12 | -0.45 | 0.000533 | NM_002969.3:425 |
| MAPK8 | 0.096 | 0.00451 | NM_002750.2:945 |
| MCM2 | -0.0885 | 0.00354 | NM_004526.2:2945 |
| MCM5 | -0.254 | 0.00695 | NM_006739.3:1580 |
| MCM7 | 0.0984 | 0.00256 | NM_182776.1:1325 |
| MED12 | -0.67 | 0.000414 | NM_005120.2:375 |
| MET | 0.0583 | 0.000913 | NM_000245.2:405 |
| MGMT | 0.149 | 0.00182 | NM_002412.3:323 |
| MLLT3 | -0.553 | 0.00496 | NM_004529.2:1480 |
| MSH6 | -0.17 | 4.64E-05 | NM_000179.1:3525 |
| MUTYH | -0.41 | 0.000465 | NM_012222.2:412 |
| NBN | -0.0622 | 0.00497 | NM_001024688.1:1105 |
| NCOR1 | 0.0934 | 0.00145 | NM_006311.3:1390 |
| NF1 | -0.062 | 0.00281 | NM_000267.2:1035 |
| NGF | 0.411 | 6.58E-05 | NM_002506.2:100 |
| NOG | -0.182 | 0.00277 | NM_005450.4:1543 |
| NOTCH2 | 0.0875 | 0.00386 | NM_024408.3:2842 |
| NPM1 | -0.216 | 0.00405 | NM_002520.5:10 |
| NRAS | -0.152 | 1.20E-05 | NM_002524.3:877 |
| NTHL1 | -0.355 | 0.000495 | NM_002528.5:476 |
| NUMBL | -0.184 | 0.00883 | NM_004756.3:591 |
| NUPR1 | 0.63 | 3.53E-05 | NM_001042483.1:829 |
| PBX1 | -0.437 | 0.000271 | NM_002585.2:368 |
| PDGFC | 0.146 | 0.00113 | NM_016205.1:10 |
| PIK3CA | -0.146 | 0.003 | NM_006218.2:2445 |
| PIK3R1 | 0.172 | 0.00185 | NM_181504.2:1105 |
| PIK3R2 | -0.125 | 0.000396 | NM_005027.2:3100 |
| PIM1 | 0.341 | 0.00226 | NM_002648.2:1630 |
| PKMYT1 | -0.135 | 0.00766 | NM_004203.3:780 |
| PLA2G3 | -0.27 | 0.00939 | NM_015715.3:2415 |
| PLA2G4C | 1.05 | 0.000181 | NM_003706.2:2310 |
| PLAT | 0.957 | 3.73E-07 | NM_000931.2:1334 |
| PLAU | -0.0725 | 0.00152 | NM_002658.2:793 |
| PLCB1 | 0.459 | 0.000508 | NM_182734.1:170 |
| PLCE1 | 0.175 | 0.0037 | NM_001165979.1:392 |
| PLD1 | 0.24 | 0.00613 | NM_002662.3:1265 |
| PML | 0.425 | 9.64E-06 | NM_002675.3:281 |

| | | | |
|---|---|---|---|
| EPHA2 | 0.0849 | 0.0104 | NM_004431.2:1525 |
| ERCC6 | 0.0841 | 0.00483 | NM_000124.2:3235 |
| ETS2 | 0.107 | 0.000263 | NM_005239.4:1175 |
| ETV4 | 0.693 | 0.00118 | NM_001079675.1:1535 |
| FANCB | 0.16 | 0.00506 | NM_152633.2:2470 |
| FANCE | -0.265 | 0.000251 | NM_021922.2:1275 |
| FANCG | -0.189 | 0.00447 | NM_004629.1:1900 |
| FAS | 0.277 | 0.00461 | NM_152876.1:1740 |
| FBXW7 | -0.136 | 0.00807 | NM_018315.4:1480 |
| FEN1 | 0.13 | 0.000721 | NM_004111.4:425 |
| FGF1 | 0.893 | 4.04E-06 | NM_033137.1:315 |
| FGF2 | -0.105 | 0.000208 | NM_002006.4:620 |
| FGFR1 | -0.177 | 0.000794 | NM_015850.2:1335 |
| FN1 | 0.436 | 8.99E-08 | NM_212482.1:1776 |
| FOS | 1.17 | 9.84E-07 | NM_005252.2:1475 |
| FOSL1 | 0.187 | 0.000545 | NM_005438.2:280 |
| FST | 0.605 | 0.00129 | NM_006350.2:575 |
| FUBP1 | -0.298 | 6.65E-05 | NM_003902.3:820 |
| FUT8 | 0.138 | 0.00723 | NM_004480.4:2841 |
| FZD2 | -0.445 | 8.59E-06 | NM_001466.2:845 |
| FZD7 | 0.427 | 0.000902 | NM_003507.1:1890 |
| GADD45A | 0.356 | 4.54E-09 | NM_001924.2:865 |
| GATA2 | 0.313 | 0.000546 | NM_032638.3:1495 |
| GNA11 | -0.0539 | 0.00818 | NM_002067.1:555 |
| GNAS | 0.0315 | 0.000164 | NM_080425.1:1910 |
| GRB2 | -0.0778 | 0.00643 | NM_002086.4:412 |
| H3F3A | -0.114 | 6.11E-05 | NM_002107.3:190 |
| H3F3C | 0.0729 | 0.00462 | NM_001013699.2:829 |
| HDAC1 | -0.0938 | 0.0031 | NM_004964.2:785 |
| HDAC10 | -0.414 | 0.00185 | NM_032019.5:932 |
| HDAC5 | -0.155 | 0.00673 | NM_005474.4:3160 |
| HDAC6 | -0.106 | 0.00736 | NM_006044.2:536 |
| HES1 | 0.573 | 2.50E-05 | NM_005524.2:860 |
| HIST1H3B | -0.36 | 1.10E-05 | NM_003537.3:335 |
| HIST1H3H | -0.768 | 5.07E-07 | NM_003536.2:355 |
| HOXA10 | -0.167 | 0.00323 | NM_018951.3:1503 |
| HSP90B1 | -0.232 | 7.47E-06 | NM_003299.1:160 |
| HSPA1A | -0.627 | 0.00078 | NM_005345.5:98 |
| HSPB1 | -0.297 | 2.80E-07 | NM_001540.3:374 |
| ID1 | -0.329 | 0.00199 | NM_002165.2:345 |
| ID2 | -0.293 | 0.000322 | NM_002166.4:505 |
| IDH1 | 0.372 | 0.000114 | NM_005896.2:105 |
| IDH2 | 0.296 | 0.000212 | NM_002168.2:944 |
| IGFBP3 | -0.13 | 0.00114 | NM_000598.4:1255 |

| | | | |
|---|---|---|---|
| POLD4 | 0.172 | 0.00716 | NM_021173.2:470 |
| POLR2H | 0.31 | 7.13E-05 | NM_001278698.1:940 |
| POLR2J | -0.385 | 0.00293 | NM_006234.4:618 |
| PPARGC1A | 0.338 | 0.000103 | NM_013261.3:1505 |
| PPP2R1A | -0.107 | 0.00886 | NM_014225.3:1440 |
| PPP3CA | -0.175 | 0.00507 | NM_000944.4:3920 |
| PPP3CC | 0.434 | 0.000341 | NM_005605.3:1460 |
| PRKACA | -0.102 | 0.000545 | NM_002730.3:400 |
| PRKAR1B | 0.234 | 0.000221 | NM_001164759.1:1112 |
| PRKAR2B | -0.205 | 0.00885 | NM_002736.2:1350 |
| PTPN11 | 0.209 | 3.37E-06 | NM_002834.3:1480 |
| PTPRR | 0.377 | 0.000247 | NM_001207015.1:1652 |
| PTTG2 | -0.1 | 0.000445 | NM_006607.2:5 |
| RAD21 | -0.112 | 3.10E-05 | NM_006265.2:1080 |
| RAD50 | 0.349 | 5.04E-07 | NM_005732.2:5397 |
| RB1 | -0.0768 | 0.0012 | NM_000321.1:2110 |
| RFC3 | 0.0397 | 0.0093 | NM_002915.3:740 |
| RHOA | -0.142 | 0.000195 | NM_001664.2:1230 |
| RPS6KA5 | 0.452 | 2.72E-05 | NM_004755.2:855 |
| RUNX1 | -0.247 | 0.000278 | NM_001754.4:635 |
| SETD2 | -0.182 | 0.00134 | NM_014159.6:6160 |
| SF3B1 | -0.0852 | 0.000348 | NM_001005526.1:0 |
| SIN3A | -0.397 | 0.000405 | NM_015477.1:1605 |
| SKP1 | -0.11 | 2.50E-05 | NM_170679.2:630 |
| SMAD2 | -0.143 | 0.000164 | NM_001003652.1:4500 |
| SMAD4 | 0.186 | 0.00874 | NM_005359.3:1370 |
| SMARCA4 | -0.153 | 0.000294 | NM_003072.3:5400 |
| SMARCB1 | -0.202 | 3.73E-06 | NM_003073.3:1060 |
| SOCS1 | 0.784 | 1.49E-05 | NM_003745.1:1025 |
| SOS1 | -0.109 | 0.00599 | NM_005633.2:1635 |
| SOS2 | 0.314 | 0.000529 | NM_006939.2:3845 |
| SPRY2 | 0.337 | 0.00054 | NM_005842.2:85 |
| SPRY4 | 0.346 | 0.0064 | NM_030964.3:1900 |
| SRSF2 | -0.202 | 0.00094 | NM_003016.3:312 |
| STAG2 | -0.153 | 0.000859 | NM_001042749.1:4040 |
| STAT1 | 0.566 | 8.49E-06 | NM_007315.2:205 |
| STK11 | -0.146 | 0.000781 | NM_000455.4:2060 |
| STMN1 | -0.175 | 0.00217 | NM_203401.1:478 |
| SUV39H2 | -0.175 | 0.00179 | NM_024670.3:2035 |
| TGFB1 | -0.219 | 1.51E-05 | NM_000660.3:1260 |
| TGFB2 | 0.274 | 2.37E-05 | NM_003238.2:1125 |
| TGFBR2 | 0.224 | 0.000309 | NM_001024847.1:1760 |
| TIAM1 | 0.831 | 0.00205 | NM_003253.2:5620 |
| TLR4 | 0.531 | 0.00021 | NM_138554.2:2570 |

| | | | |
|---|---|---|---|
| IL1RAP | 0.227 | 0.00317 | NM_002182.2:460 |
| IL6 | 0.942 | 2.98E-05 | NM_000600.1:220 |
| IL6R | 0.113 | 0.00351 | NM_000565.2:993 |
| IL7R | 0.278 | 8.56E-05 | NM_002185.2:1610 |
| IRAK2 | 0.368 | 0.00105 | NM_001570.3:1285 |
| ITGA2 | -0.213 | 0.000358 | NM_002203.2:475 |
| ITGB3 | 0.838 | 1.70E-06 | NM_000212.2:4485 |
| JAG2 | -0.291 | 0.000984 | NM_145159.1:4225 |
| JAK1 | -0.186 | 0.00128 | NM_002227.1:285 |
| JUN | 0.457 | 2.58E-06 | NM_002228.3:140 |
| KAT2B | 0.169 | 0.00135 | NM_003884.3:1220 |
| KDM6A | 0.195 | 0.00201 | NM_021140.2:2590 |
| KITLG | 0.313 | 0.000525 | NM_003994.4:1155 |
| KLF4 | 0.608 | 4.89E-06 | NM_004235.4:1980 |
| LAMA1 | 0.212 | 0.000991 | NM_005559.2:5230 |
| LAMA3 | 0.319 | 9.68E-05 | NM_000227.3:4260 |
| LAMA5 | -0.413 | 0.01 | NM_005560.3:787 |

| | | | |
|---|---|---|---|
| TNC | 0.235 | 0.000118 | NM_002160.3:4 |
| TNFAIP3 | 0.709 | 0.00011 | NM_006290.2:260 |
| TNFRSF10B | 0.436 | 4.46E-06 | NM_003842.3:565 |
| TNFRSF10D | -0.45 | 0.00692 | NM_003840.3:2380 |
| TP53 | -0.102 | 0.00227 | NM_000546.2:1330 |
| TSC1 | 0.243 | 0.00108 | NM_000368.3:100 |
| TSLP | 0.542 | 0.000111 | NM_033035.4:899 |
| TTK | -0.138 | 0.0022 | NM_003318.3:1200 |
| UBE2T | 0.12 | 0.00239 | NM_014176.3:595 |
| VEGFA | 0.622 | 4.65E-06 | NM_001025366.1:1325 |
| VHL | 0.121 | 9.16E-06 | NM_000551.2:1280 |
| WEE1 | 0.155 | 0.000653 | NM_003390.3:1225 |
| WT1 | -0.449 | 0.00213 | NM_000378.3:2160 |
| XPA | -0.182 | 0.000662 | NM_000380.3:265 |
| ZAK | -0.13 | 0.00181 | NM_016653.2:995 |
| ZIC2 | 0.447 | 0.000131 | NM_007129.2:1849 |

**Table S.3.** Significantly enriched KEGG signalling pathways of chloroquine phosphate treated liver cells (Huh7 and Mahlavu). Only the pathways that are reported to be enriched in both Enrichr[39] and Webgestalt[40] analysis are given. Significance values and scores are obtained from Enrichr.

| Cell-line | Term | Odds Ratio | Combined Score | P-value | Adjusted P-value |
|---|---|---|---|---|---|
| Huh7 | PI3K-Akt signaling pathway | 15.611 | 1352.847 | 2.31E-38 | 3.56E-36 |
| | MAPK signaling pathway | 15.165 | 1040.176 | 1.63E-30 | 1.67E-28 |
| | Cell cycle | 26.528 | 1718.160 | 7.44E-29 | 5.73E-27 |
| | Ras signaling pathway | 14.179 | 687.902 | 8.50E-22 | 2.01E-20 |
| | Apoptosis | 17.483 | 716.143 | 1.62E-18 | 2.77E-17 |
| | FoxO signaling pathway | 16.946 | 613.216 | 1.92E-16 | 2.47E-15 |
| | TGF-beta signaling pathway | 19.006 | 559.205 | 1.67E-13 | 1.83E-12 |
| | Th17 cell differentiation | 15.986 | 433.755 | 1.65E-12 | 1.54E-11 |
| | p53 signaling pathway | 20.102 | 516.678 | 6.88E-12 | 5.05E-11 |
| | Longevity regulating pathway | 15.480 | 383.006 | 1.80E-11 | 1.18E-10 |
| | TNF signaling pathway | 14.354 | 342.161 | 4.44E-11 | 2.74E-10 |
| | T cell receptor signaling pathway | 14.330 | 314.235 | 3.00E-10 | 1.74E-09 |
| | Gap junction | 14.952 | 305.973 | 1.30E-09 | 7.01E-09 |
| | Toll-like receptor signaling pathway | 8.856 | 98.595 | 1.46E-05 | 4.33E-05 |
| | cGMP-PKG signaling pathway | 6.341 | 64.137 | 4.05E-05 | 1.13E-04 |
| | Cytokine-cytokine receptor interaction | 3.580 | 22.484 | 0.002 | 0.004 |
| Mahlavu | MAPK signaling pathway | 14.860 | 1427.261 | 1.93E-42 | 2.97E-40 |
| | PI3K-Akt signaling pathway | 12.641 | 1137.673 | 8.20E-40 | 6.31E-38 |
| | Ras signaling pathway | 13.384 | 855.224 | 1.77E-28 | 2.73E-27 |
| | Focal adhesion | 11.932 | 547.317 | 1.20E-20 | 1.09E-19 |
| | JAK-STAT signaling pathway | 12.402 | 493.162 | 5.38E-18 | 3.60E-17 |
| | TNF signaling pathway | 15.774 | 617.099 | 1.02E-17 | 6.71E-17 |
| | Apoptosis | 12.134 | 413.198 | 1.63E-15 | 9.10E-15 |
| | HIF-1 signaling pathway | 14.612 | 465.393 | 1.47E-14 | 7.43E-14 |
| | TGF-beta signaling pathway | 15.221 | 465.149 | 5.34E-14 | 2.49E-13 |
| | Toll-like receptor signaling pathway | 13.172 | 373.438 | 4.87E-13 | 2.14E-12 |
| | Oxytocin signaling pathway | 8.953 | 203.071 | 1.41E-10 | 5.30E-10 |
| | Cholinergic synapse | 8.969 | 152.336 | 4.21E-08 | 1.32E-07 |
| | IL-17 signaling pathway | 9.820 | 161.257 | 7.38E-08 | 2.25E-07 |
| | NOD-like receptor signaling pathway | 6.670 | 108.520 | 8.59E-08 | 2.59E-07 |
| | Necroptosis | 5.637 | 63.755 | 1.23E-05 | 3.07E-05 |
| | Circadian entrainment | 5.649 | 41.080 | 0.001 | 0.001 |
| | Renin secretion | 6.618 | 46.006 | 0.001 | 0.002 |
| | Endocytosis | 2.994 | 15.531 | 0.006 | 0.011 |

**Table S.4.** The list of genes/proteins shared between the CROssBAR COVID-19 knowledge graph and the differentially expressed genes of the chloroquine phosphate treated liver cells (Huh7 and Mahlavu cell-lines) found using the NanoString platform.

| Gene name | Protein accession | Protein name |
|---|---|---|
| STAT1 | P42224 | Signal transducer and activator of transcription 1-alpha/beta |
| VHL | P40337 | von Hippel-Lindau disease tumor suppressor |
| GNAS | Q5JWF2 | Guanine nucleotide-binding protein G(s) subunit alpha isoforms Xlas |
| MAPK12 | P53778 | Mitogen-activated protein kinase 12 |
| PRKACA | P17612 | cAMP-dependent protein kinase catalytic subunit alpha |
| XPA | P23025 | DNA repair protein complementing XP-A cells |
| BCL2L1 | Q07817 | Bcl-2-like protein 1 |
| CTNNB1 | P35222 | Catenin beta-1 (Beta-catenin) |
| B2M | P61769 | Beta-2-microglobulin |
| CHEK2 | O96017 | Serine/threonine-protein kinase Chk2 |
| BAP1 | Q92560 | Ubiquitin carboxyl-terminal hydrolase BAP1 |
| PRKAR2B | P31323 | cAMP-dependent protein kinase type II-beta regulatory subunit |
| PPP2R1A | P30153 | Serine/threonine-protein phosphatase 2A 65 kDa regulatory subunit A alpha |
| RFC3 | P40938 | Replication factor C subunit 3 |
| DNMT1 | P26358 | DNA (cytosine-5)-methyltransferase 1 |
| MNAT1 | P51948 | CDK-activating kinase assembly factor MAT1 |
| DAXX | Q9UER7 | Death domain-associated protein 6 |

**Table S.5.** CROssBAR COVID-19 large-scale and simplified knowledge graphs node and edge statistics.

| **Large-scale COVID-19 KG** | |
|---|---|
| **Node_Type** | **Node_Stat** |
| Human proteins | 475 |
| SARS-CoV-1 proteins | 33 |
| SARS-CoV-2 proteins | 31 |
| Drugs | 108 |
| Compounds | 233 |
| KEGG Pathways | 23 |
| Reactome Pathways | 34 |
| HPOs | 27 |
| KEGG Diseases | 10 |
| EFO Diseases | 13 |
| Organism | - |
| **TOTAL** | **987** |
| **Edge_Type** | **Edge_Stat** |
| PPIs | 1,284 |
| Approved/investigational DTIs* | 135 |
| Bioassay-based exp. DTIs (ChEMBL) | 335 |
| Predicted DTIs | 326 |
| Protein-KEGG Pathway associations | 244 |
| Protein-Reactome Pathway associations | 474 |
| Protein-KEGG Disease associations | 31 |
| Protein-EFO Disease associations | 21 |
| Protein-HPO associations | 653 |
| HPO-EFO Disease associations | 41 |
| Drug-KEGG Disease Indications | 1 |
| Drug-COVID-19 Disease Indications | 29 |
| KEGG Pathway-Disease modulations | 2 |
| Orthology relations between SARS-CoVs | 29 |
| Protein complex - subunit relations (virus) | 34 |
| **TOTAL** | **3,639** |

| **Simplified COVID-19 KG** | |
|---|---|
| **Node_Type** | **Node_Stat** |
| Human proteins | 16 |
| SARS-CoV-1 proteins | 15 |
| SARS-CoV-2 proteins | 14 |
| Drugs | 46 |
| Compounds | 36 |
| KEGG Pathways | 12 |
| Reactome Pathways | 7 |
| HPOs | 18 |
| KEGG Diseases | 4 |
| EFO Diseases | 7 |
| Organism | 3 |
| **TOTAL** | **178** |
| **Edge_Type** | **Edge_Stat** |
| PPIs | 45 |
| Approved/investigational DTIs | 12 |
| Bioassay-based exp. DTIs (ChEMBL) | 37 |
| Predicted DTIs | 12 |
| Protein-KEGG Pathway associations | 26 |
| Protein-Reactome Pathway associations | 15 |
| Protein-KEGG Disease associations | 4 |
| Protein-EFO Disease associations | 5 |
| Protein-HPO associations | 12 |
| HPO-EFO Disease associations | 30 |
| Drug-KEGG Disease Indications | 6 |
| Drug-COVID-19 Disease Indications | 29 |
| KEGG Pathway-Disease modulations | 8 |
| Orthology relations between SARS-CoVs | 12 |
| Organism-gene/protein relations | 45 |
| **TOTAL** | **298** |

*DTIs: drug/compound-target interactions

**Table S.6.** Literature based information for new potential COVID-19 based repurposing of CROssBAR COVID-19 knowledge graph drugs.

| **Drug Name** | **DrugBank id** | **Description** | **Source **** | **Clinical trial Id** | **Current state *** |
|---|---|---|---|---|---|
| Cyclosporine | DB00091 | calcineurin inhibitor | Enrichment (DrugBank & ChEMBL) | NCT04392531 | Phase 4 |
| Tocilizumab | DB06273 | IL-6 inhibitor | Enrichment (DrugBank) | NCT04377750 | Phase 4 |
| Amlodipine | DB00381 | calcium channel blocker | DL Prediction (MDeePred) | NCT04330300 | Phase 4 |
| Siltuximab | DB09036 | IL-6 inhibitor | Enrichment (DrugBank) | NCT04330638 | Phase 3 |
| Prednisolone | DB00860 | glucocorticoid steroid | Enrichment (ChEMBL) | NCT04381936 | Phase 2-3 |
| Vazegepant | DB15688 | calcitonin gene-related peptide (CGRP) receptor antagonist | DL Prediction (DEEPScreen) | NCT04346615 | Phase 2-3 |
| Quercetin | DB04216 | polyphenolic flavonoid | Enrichment (DrugBank) | NCT04377789 | Not Applicable |
| Artenimol | DB11638 | artemisinin derivative and antimalarial agent | Enrichment (DrugBank) | - | *In silico* study |
| Lifitegrast | DB11611 | integrin antagonist | Enrichment (DrugBank) | - | *In silico* study |
| Amcinonide | DB00288 | corticosteroid | DL Prediction (MDeePred) | - | *In silico* study |
| Becatecarin | DB06362 | diethylaminoethyl analogue of rebeccamycin | DL Prediction (MDeePred) | - | *In silico* study |
| Quinfamide | DB12780 | antiprotozoal agent | DL Prediction (MDeePred) | - | *In silico* study |
| Rocaglamide | DB15495 | eIF4A inhibitor | Enrichment (DrugBank) | - | - |
| Didesmethyl rocaglamide | DB15496 | eIF4A inhibitor | Enrichment (DrugBank) | - | - |

* Some of these drugs have multiple clinical trials concerning COVID-19. In these cases, the one with the latest phase is given.
** The source of the drug prediction as either enrichment analysis or DL prediction (prediction of our deep learning-based tools).
